# Exploring the probiotic potential, antioxidant capacity, and healthy aging based on whole genome analysis of *Lactiplantibacillus plantarum* LPJBC5 isolated from fermented milk product

**DOI:** 10.1101/2024.03.14.584937

**Authors:** Anupam Bhattacharya, Tulsi K. Joishy, Mojibur R. Khan

**Author notes:** Corresponding author: Prof. M. R. Khan, Life Sciences Division, Institute of Advanced Study in Science and Technology (IASST), Guwahati-781035, Assam, India. Contributed equally (Anupam Bhattacharya, Tulsi K. Joishy).

## Abstract

*Lactiplantibacillus plantarum* is a beneficial bacterium commonly found in fermented foods, including fermented milk products. In the present study, we reported the whole genome sequence of *L. plantarum* LPJBC5. The complete genome sequence of LPJBC5 was 3.23Mb, and the average GC% was found to be 44.55% encoding a total of 3016 genes. A comprehensive analysis of the LPJBC5 genome detected major carbohydrate-active enzymes, exopolysaccharide synthesis genes (*eps*), and the presence of genes related to stress response and antioxidant activity. Longevity regulating genes including *kat*E, CAT (chloramphenicol acetyltransferase), *cat*B and their regulating pathways (MAPK signaling pathway) were detected in the LPJBC5 genome. The genome was found to contain plantaracin (*pln)* operon for the production of antimicrobial bacteriocin and various regions of secondary metabolite biosynthetic gene clusters, including the type III polyketide synthases (T3PKS), Ribosomally synthesized and post-translationally modified peptide product (Ripp-Like), which exhibit specific antimicrobial activity. A comparative pangenome analysis was performed to further evaluate the metabolic framework of L. plantarum JBC5, including the complete *L. plantarum* genome (N=30) retrieved from publicly available repositories. The core/soft-core genome was found to harbor probiotic associated marker genes. Functional analysis revealed the presence of genes majorly enriched in cell wall/membrane/envelop biogenesis, carbohydrate metabolism and transport, amino acid metabolism and transport, translational mechanism, and transcription processes. Identification of probiotic and longevity associated genes suggests that LPJBC5 holds potential as a probiotic candidate with multifaceted applications in the food industries.

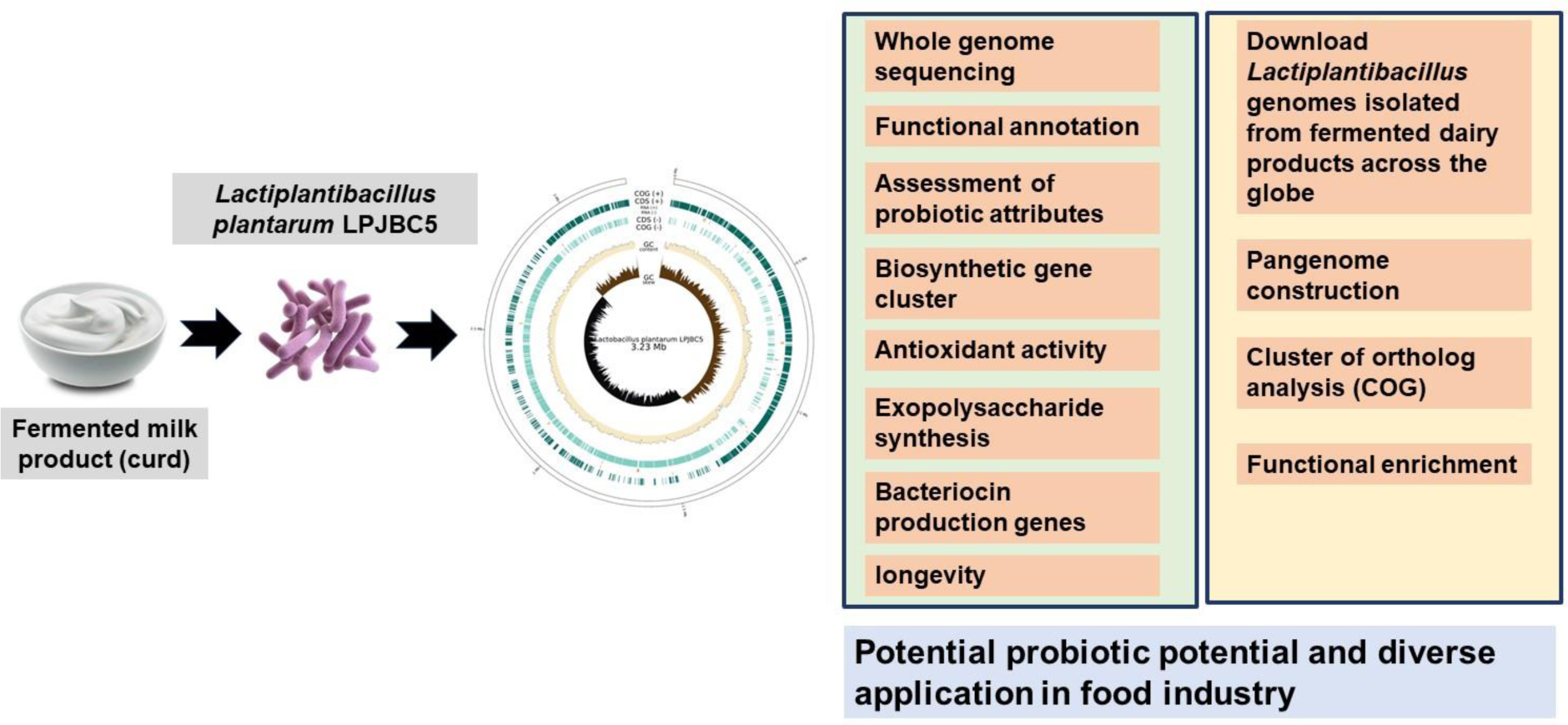

## Introduction

*Lactiplantibacillus plantarum* is majorly found in fermented food and functions as a probiotic. It carries one of the largest genomes among the lactic acid bacteria (LAB) (Abedin, Chourasia et al. 2023). The *L. plantarum* is considered one of the most crucial probiotic employed in food industry for the development of functional food (Behera, Ray et al. 2018, Damián, Cortes-Perez et al. 2022, Abedin, Chourasia et al. 2023). *L. plantarum* strains were isolated from various fermented food products and exhibit beneficial effects including, gut health (Rastogi and Singh et al. 2022), maintenance of gut microbiome homeostasis (Qiu, Tao et al. 2018), exert potent anti-obesity effects in mice (Soundharrajan, Kuppusamy et al. 2020), prevention of various diseases including inflammatory bowel disease (Duary et al. 2012), liver damage (Choi, Lee et al. 2019), uremia (Patra, Mandal et al. 2018). The underlying molecular mechanism exhibiting these probiotic properties from a genomic perspective remains unclear. Several studies were conducted on genomic(Zhang, Lv et al. 2018, Goel, Halami et al. 2020) and pan-genome analysis (Choi, Jin et al. 2018, Carpi, Coman et al. 2022) of *L. plantarum* strains based on publicly available genomic datasets.

The report suggests the beneficial effects are strain-specific (Ramos, Thorsen et al. 2013). Exploring LAB species in fermented food may reveal novel strains with functional properties. In our previous studies, fermented milk product (curd) was prepared using boiled milk and raw milk from dairy farms in India, and a potential probiotic bacterium, L. plantarum, was isolated and characterized. The strain LPJBC5 promoted longevity and diverse aspects of healthy aging, including delayed age-related physical functions in C. elegans.(Joishy, Dehingia et al. 2019, Kumar, Joishy et al. 2022). In the present study, a whole genome of *L. plantarum* LPJBC5, which was isolated from fermented milk product, was sequenced. Functional genomic analysis was performed for general genomic characteristics, identification of strain-specific bacterial genes responsible for longevity, antioxidant properties, adhesions, production of exopolysaccharides, and secondary metabolites biosynthetic gene clusters. A comprehensive pan-genome analysis of publicly available *L. plantarum* genomes isolated from fermented milk products was conducted to gain insight into the unique and shared genes involved in various biochemical and metabolic pathways by different strains. Distribution of clusters of orthologous genes, KEGG pathways of core, accessory, and unique genes *L. plantarum* pan genomes were also studied. This study may provide useful genomic insight into the various properties of *L. plantarum* LPJBC5, including healthy aging, and make it a potential probiotic candidate in the food industry.

## Material and Methods

### Bacterial growth condition and DNA extraction

*Lactiplantibacillus plantarum* JBC5 was grown in MRS medium (Himedia, India) at 30°C. The genomic DNA was extracted from the overnight grown culture using GenElute™ Bacterial Genomic DNA Kit (St. Louis, USA).

### Whole genome sequencing and assembly of *L. plantarum* LPJBC5

Whole-genome sequencing libraries were prepared using a KAPA DNA HyperPrep kit as per instruction. In brief, the DNA is sheared using a Covaris ultrasonicator. Sheared DNA is subjected to a sequence of enzymatic steps for repairing the ends and tailing with dA-tail followed by ligating indexed adapter sequences. These adapter ligated fragments are then cleaned up using SPRI beads. Next, the clean fragments are indexed using limited cycle PCR to enrich the adapter ligated molecules. The amplified products are purified and checked for quality and quantity before sequencing. Prepared libraries sequenced on Illumina Novaseq platform to generate 2x150 bp reads/sample. Sequenced data processed to generate FASTQ files. The quality of the reads was checked based on quality score distribution, average base content per read, GC distribution, PCR amplification issue, overrepresented sequences, and adapters. Based on the quality inspection, sequences were trimmed to retain high-quality reads for further analysis. The adapter was removed using fastq mcf (v- 1.04.803). The processed reads were aligned to the reference genome of bacteria, viruses and archaea to check the alignment percentage using Burrows-Wheeler Aligner (BWA) (version 0.7.12) (Li and Durbin et al. 2009). De-novo assembly was performed using a megahit assembler (v1.2)(Li, Liu et al. 2015).a Megahit uses succinct de Bruijn graphs which are compressed representations of de Bruijn graphs. The assembled scaffolds were taken for further downstream analysis. The assembled sequence was screened for ribosomal genes (*rps*) for identification using rMLST approach (Jolley, Bliss et al. 2012).

### Gene prediction and functional annotation of *L. plantarum* LPJBC5 genome

The genome was annotated by using Prokaryotic Genome Automatic Annotation Pipeline (PGAP) (Zhao, Wu et al. 2012), RASTk (Brettin, Davis et al. 2015), Prokka (Seemann et al. 2014) to get a comprehensive insight into the list of genes present in the genome of *L. plantarum* LPJBC5. Briefly, ARAGORN(Laslett and Canback et al. 2004), Prodigal(Hyatt, Chen et al. 2010) and GeneMarkS-2+(Lomsadze, Gemayel et al. 2018) annotated the genes related to tRNA, rRNA and protein coding sequences. The protein coding genes were further classified based on domain-based search through HMMER, Pfam (Potter, Luciani et al. 2018, Mistry, Chuguransky et al. 2021), and TIGRFAMs (Haft, Loftus et al. 2001) database. Annotated genes were compared against a cluster of orthologous groups. The orthologous groups were functionally annotated using eggNOG (Huerta-Cepas, Szklarczyk et al. 2019), KEGG pathways, and SMART & PFAM domains to gain more insight into complex biological functions and pathways. BlastKOALA (Kanehisa, Sato et al. 2016) was used to annotate genes based on the KEGG database. The genes were functionally curated based on probiotic attributes, including Stress response, adhesion, acid tolerance, and bile resistance. The annotated genome was also tested for probiotic potential using iProbiotics (Sun, Li et al. 2021).

### Identification of secondary metabolites and biosynthetic gene clusters

Antismash (Medema, Blin et al. 2011) was used to detect secondary metabolites biosynthesis gene clusters in the genome. The pipeline involves the identification of gene clusters based on signatures of secondary metabolites and comparative analysis using secondary metabolism clusters of orthologous groups. The genome was screened for the production of bacteriocins through a knowledge-based database in BAGEL(van Heel, de Jong et al. 2018). The program ranked the features and identified sets of putative bacteriocin gene clusters in the genome. The *L. plantarum* LPJBC5 genome was screened for antimicrobial resistance genes in the Comprehensive Antibiotic Resistance Database (CARD)(McArthur, Waglechner et al. 2013).

### Pangenome analysis of *L. plantarum* LPJBC5 and phylogenetic reconstruction of *L. plantarum* strains

A total of 30 publicly available genome datasets belonging to the *L. plantarum* species, isolated from dairy products and *L. plantarum* type strains (WCSF1 and ATCC14917), were comprehensively studied for pangenome analysis. The genomes were obtained from India, Ireland, the United Kingdom, The Netherlands, Croatia, China, Malaysia, South Africa, Italy, Slovenia, Spain, and Croatia. (**Supplementary file S1)**. Prokka (version 1.14.5) pipeline was used to annotate the genome assemblies. Pangenome profile was created in the Roary and BPGA pipeline (Page, Cummins et al. 2015, Chaudhari, Gupta et al. 2016) to identify core and accessory genes shared by *L. plantarum* strains. Four different classes of genes, including those belonging to the core (strains between 99% and 100%), soft-core (strains between 95% and 99%), shell (strains between 15% and 95%), and cloud (strains between 0% and 15%) groups, were obtained. The core and accessory genomes were aligned to construct core and pan-phylogenetic tree. Phandango (Hadfield, Croucher et al. 2017) was used to visualize the pangenome. Further to identify the functional perspective of the genes, orthologous clusters were determined using the orthoMCL algorithm (Li, Stoeckert et al. 2003).

### Functional enrichment analysis of the core and accessory genes

To identify the orthologous clusters, orthoMCL algorithm was used (Li, Stoeckert et al. 2003). To gain insights into the biological roles and potential metabolic pathways of the identified genes several databases were used including gene ontology (GO) and Kyoto Encyclopedia of Genes and Genomes (KEGG). The functional enrichment of the core and accessory genes were conducted based on biological process libraries, molecular function libraries. The statistically significant gene ontology (GO) terms (*p*-value≤ 0.05) were further considered for downstream analysis. Mobile genetic elements (MGEs) present in the genome of *L. plantarum* were detected through MobileElementFinder (Johansson, Bortolaia et al. 2021).

## Results

### Genomic features of *L. plantarum* LPJBC5

The complete genome of *L. plantarum* LPJBC5 was sequenced in the Illumina Novaseq platform, and raw data were generated. A total of 15431204 reads were generated. After trimming the adapters, 99.14% of the reads were retailed. The genome size is 3.23Mb, and the average GC% was found to be 44.55%. A total of 3016 genes, 16 ribosomal RNA genes, and 65 transfer RNA genes were present in the genome. The annotation pipeline revealed 3016 coding sequences, including 1749 and 1268 sequences assigned as functional and hypothetical proteins, respectively (**Figure 1).**

**Figure 1.**
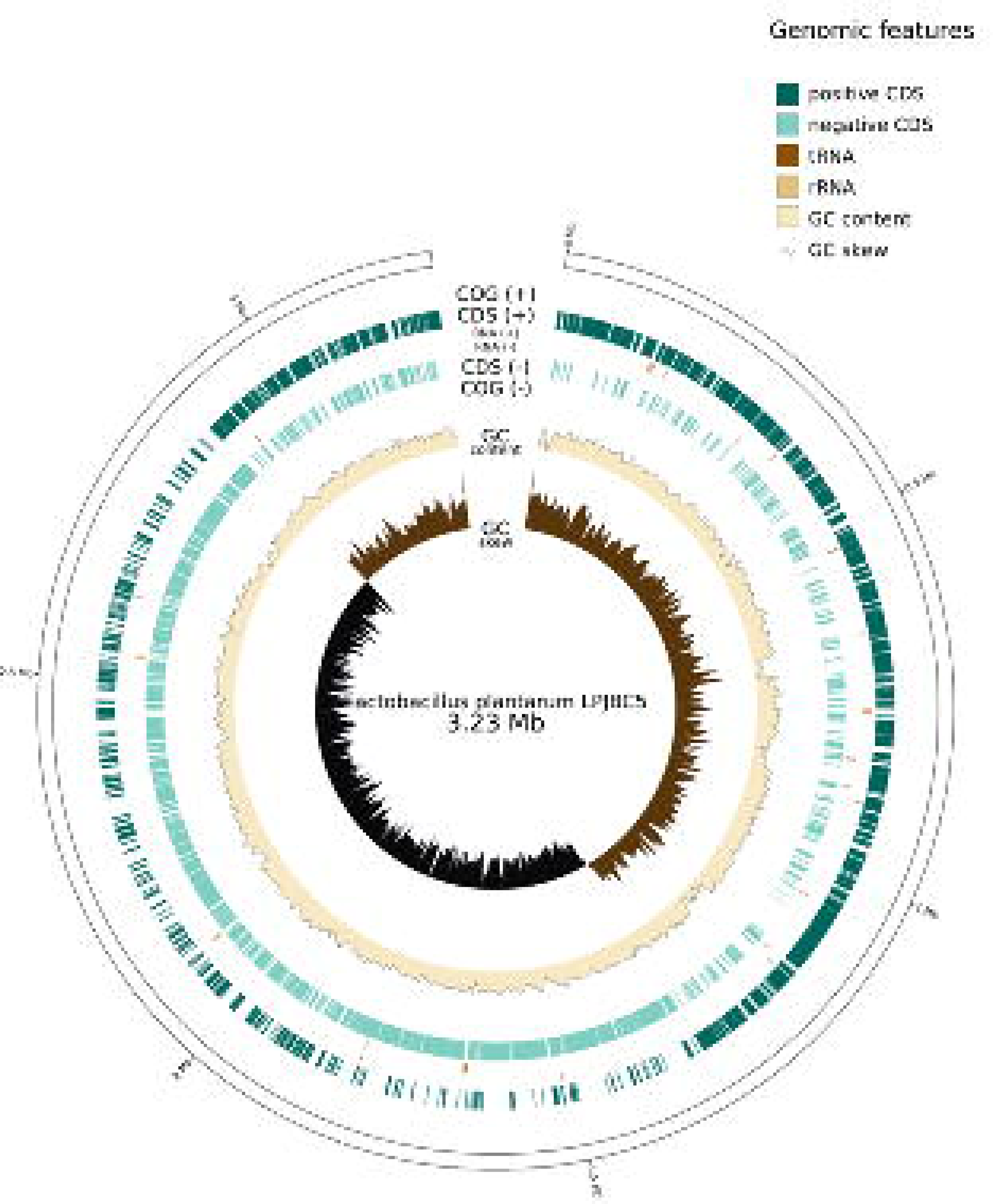
Genetic map of *L. plantarum* LPJBC5 genome. *L. plantarum* LPJBC5 genome is drawn in a single circle. Positive and negative strand coding sequences (CDSs), GC content, and GC skew are displayed.

### Characterization of bacterial strain based on whole genome sequencing

We used the Multilocus sequence typing (MLST) method for precise characterization of the L. plantarum LPJBC5 genome using the sequences of internal fragments of housekeeping genes. The whole genome was screened for conserved ribosomal genes by using rMLST approach against the BIGSdb database. Fifty-three genes encoding the bacterial ribosome protein subunits (*rps* genes) to integrate microbial taxonomy and typing were taken into account. The result revealed that the isolated strain belong to *L. plantarum*. Whole genome based blast analysis showed a significant similarity of genes and domains in the LPJBC5 genome with existing *L. plantarum* genomes (**Supplementary file S2**). Additionally, we compared the LPJBC5 genome with the existing *L. plantarum* genomes, and average nucleotide identity (ANI) values ranged from 99.42%-99.74% (**Supplementary Table S1)**.

### 2.1 Assessment of probiotic characteristics of *L. plantarum* LPJBC5

#### Analysis of stress-related genes

In the genome of *L. plantarum* LPJBC5, many adaptive stress response genes were identified, including universal stress response genes (*usp*) and proteases involved in heat stress reactions. The genes associated in stress response include cold shock-domain family proteins (*csp*A, *csp*C), Clp protein family (*clp*P,*clp*C, *clp*E, *clp*B, *clp*X), small heat shock proteins (sHSP), heat shock proteins HSP20 family (HSP33, HSP 70, HSP20, HSP2, HSP3, HSP1), *grp*E, *htp*X, and heat shock protein 10 kDa family chaperone (GroES, GroEL). Additionally, *exo*A, *lux*S gene, was also detected which are involved in stress response. For the persistent osmotic stress in the gastrointestinal tract, LPJBC5 genome encoded a multi-component binding protein dependent transport system (*opuABCD*), that assists in the osmotic environment in response to external osmotic pressure. Additionally, *gla_2, glpF_1, glpF_2* genes were also identified in the genome. The genome also contains genes involved in oxidative stress (*msr*B, *npr, nfr*A, *trx*A, *trx*B).

#### Adhesions factors present in the genome of *L. plantarum* LPJBC5

The presence of genes involved in adhesion to the gastrointestinal tract in the genome of *L. plantarum* LPJBC5 was studied as adhesion factors believed to be involved in helping probiotic strains to persist in the host gastrointestinal tract. LPJBC5 genome encodes genes involved in cell adhesion, including mucin binding domain (MucBP), fibronectin-binding proteins, collagen-binding domain, mucus adhesion promoting protein genes (mapA), enolases, LPXTG anchored proteins, and MBG domain. It was observed that the genome contains *a YceG type domain, an endolytic murein transglycosylase, with genes (mltG, yceG) associated with* adhesion under heat stress.

#### Acid and Bile resistance

The *L. Plantarum* LPJBC5 genome encoded genes associated with acid tolerance (*ldh, clpP_1, pls*C*, rel*A*, pyk, clp*B*, gpmb, gua*A*, yjb*M*, gro*L*, lux*X*, uvr*A*, tpi*A*, rec*A, ATP synthase subunit, *ywa*C, *gad*B, *pgi*), bile resistance (*cbh, opp*A*, glf, dps,lpd*A*, pyr*G*, rps*E*, rpl*F*,rpl*E*,rps*C*,rpl*D*, arg*S*, nag*B*, pep*O*, gln*A*, oppA_1, oppA_2*), acid/bile resistance (*cop*A*, arc*B*, dna*K,*dna*J, *grp*E, gro*S*, *pgk*, *eno*), intestinal persistence (*tre*A*, cel*B).

### 2.2 **Genes involved in the synthesis of exopolysaccharide**

The functional annotation revealed genes involved in the synthesis of exopolysaccharides. *L. plantarum* LPJBC5 encoded *eps* gene cluster including *eps*C, *eps*D, *eps*F, *eps*H, *eps*B. Mur ligases play an essential role in the biosynthesis of bacterial cell-wall peptidoglycan, thus represent attractive targets for the design of novel antibacterials. We investigated the presence of these genes in the genome and found the presence of *mur*B, *mur*C, *mur*D, *mur*E, *mur*F, *mur*G *mur*T in Mur ligase family, which catalyzes the stepwise formation of the peptide moiety of the peptidoglycan disaccharide peptide monomer unit. Additionally, several genes responsible for the synthesis of exopolysaccharides were distributed throughout the genome (**Table 2**).

**Table 1.** Genes related to probiotic characteristics present in *L. plantarum* LPJBC5 genome.

| Category | Gene/Gene family |
| --- | --- |
| Stress response | Cold shock-domain family proteins (*cspA*, *cspC*), Clp protein family (*clpP*, *clp_1*, *clp_2*, *clpC*, *clpE*, *clpB*, *clpX*), small heat shock proteins(sHSP), HSP33, HSP70, HSP20, HSP2, HSP3, heat shock proteins( HSP20 family, *grpE*, *HtpX*), Heat shock protein (*GroES*, *GroEL*), Universal stress protein (*usp*) |
| Adhesion | mucin binding domain(MucBP), fibronectin binding protein, fibronectin/fibrinogen binding protein, Sortase (*strA*), *exosortase*, mucus adhesion promoting protein genes (*mapA*), enolases, LPXTG anchored proteins, *exoA*, *lspA*, *pycA* |
| Immunomodulation | *dltB*, *dltD* |
| Bile resistance | *cbh*, *oppA*, *oppA_3*, *glf*, *dps*, *lpdA*, *pyrG*, *rpsE*, *rplF*, *rplE*, *rpsC*, *rplD*, *argS*, *nagB*, *pepO*, *glnA*, *oppA_1*, *oppA_2*, *ponA* |
| **Osmotic stress** | *gla_2, glpF_1, glpF_2, opuCB, opuCA, opuCD, opuAC, osmV* |
| **Acid tolerance** | *ldh, ldh_1_c, ldh_1_N, ldh_1, ldh_2, ldh_3, clpP_1, clpP_2, clpP, plsC, relA, pyk, clpB, gpmb, guaA, yjbM, groL, uvrA, uvrA_1, uvrA_2, uvrA_3, tpiA, recA, ATP synthase subunit, ywaC, gadB, pgi, plsC, dltA, luxS, atpB atpE, atpF, atpH, atpA, atpG, atpD, atpC* |
| **Acid stress/Bile resistance** | *copA, arcB, dnaK, dnaJ, grpE, groS, pgk, eno_1, eno_2, eno, htrA* |
| **Oxidative stress** | *msrA, msrB, npr, nfrA, trxA, trxB* |
| **Gut persistence** | *treA, celB* |

**Table 2.** Genes of *L. plantarum* LPJBC5 that encode proteins, involved in biosynthesis of exopolysaccharide components.

| Category | Genes |
| --- | --- |
| Synthesis of exopolysaccharide | ***eps* gene cluster**<br>*epsC, epsD, epsF, epsH, epsB*<br>***Mur* ligase gene family**<br>*murB, murC, murD, murE, murF, murG murT*<br>**others genes involved**<br>*lysA, wzx, tagA, pbp, wzy* |

### 2.3 **Genes associated with antioxidant activity**

Genomic analysis revealed the presence of NADH oxidase, NADH peroxidase, catalase, glutathione reductase, glutathione peroxidase, thioredoxin and thioredoxin reductase in *L. plantarum* LPJBC5 genome. The genome encoded various nicotinamide adenine dinucleotide (NADH) oxidation-related genes, including *npr* genes encoding NADH peroxidase. *L. plantarum* LPJBC5 genome harbored the genes related to thioredoxin system including *trx*A (encoding thioredoxin), *trx*B (encoding thioredoxin reductase), and *tpx (*encoding thiol peroxidase). The genome also contains *nrd*H (encoding glutaredoxin).

### 2.4 Genes associated with longevity in *L. plantarum* LPJBC5 genome

The genomic region of LPJBC5 was annotated to perform KO (KEGG Orthology) assignments to characterize individual gene functions. BRITE hierarchies and KEGG modules were mapped to infer high-level functions of the organism. The KEGG pathway modules revealed longevity regulating pathway (KO04211), longevity-regulating pathway-worm (KO04212), longevity-regulating pathway-multiple species (KO04213), MAPK signaling pathway (KO04011). All the above-mentioned pathways were associated with aging (KO09149). Further, an in-depth analysis revealed that several genes including catalase, *kat*E, CAT (chloramphenicol acetyltransferase), *cat*B, *srp*A were found in the LPJBC5 genome (**Supplementary figure S1**).

### 2.5 Analysis of Carbohydrate-Active EnZymes (CAZymes) in *L. plantarum* LPJBC5 genome

The analysis of CaZymes showed a total number of 198 genes present in the genome of *L. plantarum* LPJBC5 distributed in five CaZymes gene families: 52 glycoside hydrolase (GH) genes, 37 glycosyl transferase (GT) genes, 4 carbohydrate esterase (CE) genes, 12 carbohydrate-binding modules (CBM), 2 auxiliary activity (AA) genes (**Figure 2)**. 39 CaZymes gene clusters (CGCs) were identified that contain CaZymes, Transcription factors (TF), and transporters (TCs) to synergistically degrade/synthesize various highly complex carbohydrates. Analysis revealed that these gene clusters utilized various substrates including starch, glycogen, cellobiose, alpha-mannan, glycogen, galactan, and beta-glucoside (**Supplementary file S3).**

**Figure 2.**
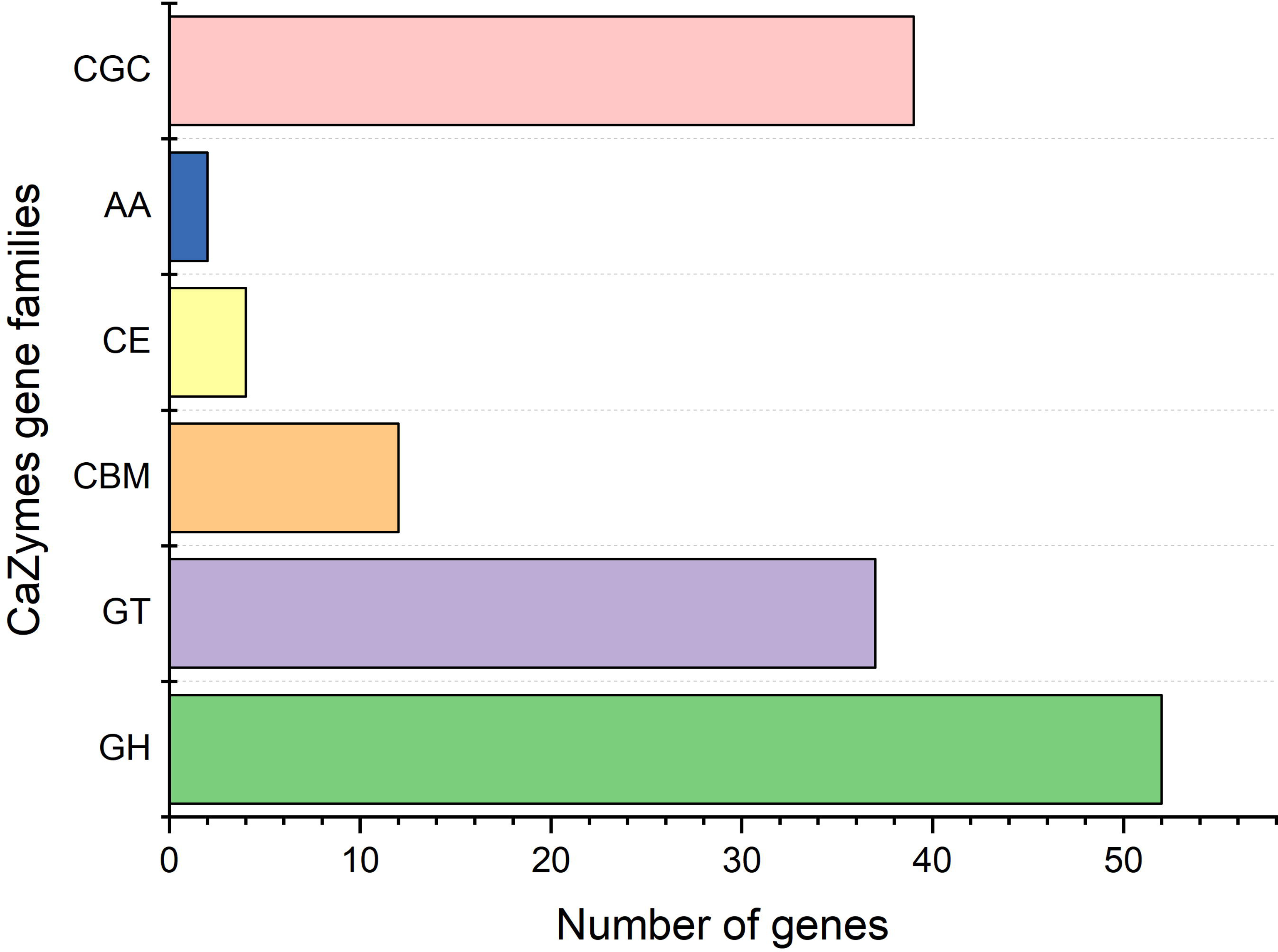
The figure depicts the genes and gene clusters involved in different category of carbohydrate active enzymes present in *L. plantarum* LPJBC5.

### 2.6 **Biosynthetic gene cluster detected in the genome**

Genomic data mining identified many of the gene clusters coding for bioactive secondary metabolites, including terpene, T3PKS, Ribosomally synthesized and post-translationally modified peptide product (Ripp-Like), and cyclic-lactone-autoinducer. The results revealed the detected biosynthetic gene clusters (Ripp-like) located in 24991 to 27141 bp region in *L. plantarum* LPJBC5 genome, which showed significant similarity against other *L. plantarum* genome (NZ_CP016270, NZ_CP019722, NZ_CP016071, NZ_CP017354, NZ_CP053337, NZ_CP013130, NZ_CP048921, NZ_CP025988, NZ_CP032744, NZ_CP059294) ranging from 99% identity with an e-value of 0.0 (Supplementary figure 5). The T3PKS region spans 1503836bp - 1505005 bp in the genome. While comparing with other *L. plantarum* strains (NZ_CP026743, NZ_CP025991, NC_004567, NZ_CP012343, NZ_CP028229, NZ_CP017954, NZ_CP050805, NZ_CP046935, NZ_CP030105, NC_021514), ranging from 97% to 100% sequence similarity was observed against this cluster (supplementary figure 6). Terpene domain is located in 2,497366bp - 2498247 bp region of the genome and found 100% similarity with existing *L. plantarum* genomes (NZ_CP020816, NZ_CP023771, NZ_MKEG01000007, NZ_P053571, NZ_CP017354, NZ_CP026743, NZ_CP023728, NZ_CP059294, NZ_CP035223, NZ_CP021528) (**Supplementary figure S2**).

### 2.7 Putative bacteriocin-related genes identified in *L. plantarum* LPJBC5

Putative bacteriocin operon in the genome was screened, and different classes of Bacteriocins were identified. Significant similarly was observed with Plantaricin family, which includes plantaricinK (Evalue=8.68e-39 match=100.00%), plantaricinJ (evalue=3.37e-38 match=100.00%;), plantaricinN (evalue=1.41e-36 match=100.00%), plantaricinA (evalue=1.82e-30 match=100.00%), plantaricinF (evalue=3.59e-36 match=100.00%), PlantaricinE (evalue=3.77e-38 match=100.00%). Additionally, *pln*O (evalue=0.0, match=99.252%), *pln*C (evalue=4.09e-163, match=99.550%), *pln*D (evalue=0.0, match=99.190%,), *pln*S (evalue=3.66e-09, match=38.889%), *pln*L (evalue=0.0, match=100.000%) were detected in the genome. The identified open reading frames, along with their positions, corresponding sequences, percent identity, and annotations, were provided in the **Figure 3 and Supplementary file S4.**

**Figure 3.**
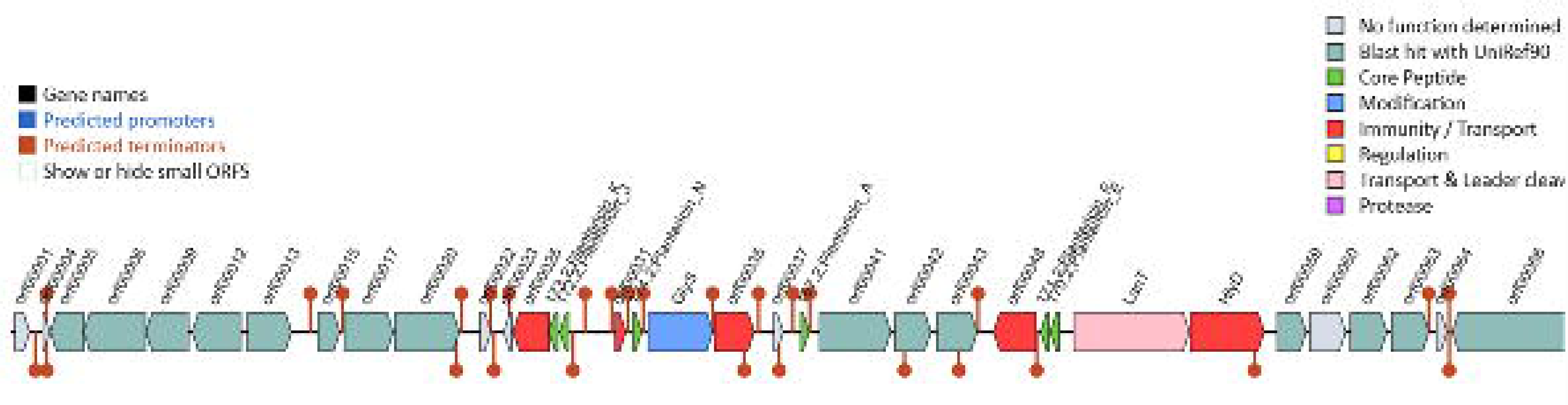
Putative bacteriocin ORFs (antimicrobial peptides) based on a database containing information about known bacteriocins and adjacent genes involved in bacteriocin activity.

### 2.8 **Antimicrobial resistance genes**

The genome was screened for potential antimicrobial resistance genes (AMR) and detected glycopeptide resistance gene cluster, tetracycline resistance genes [tet(A), tet(B),tet(R), tet(O)], major facilitator superfamily (arlR, bmr, mdtG), ATP-binding cassette (macB, bcrA). The complete lists of potential AMR genes, including high and low stringency filter, drug class, and resistance mechanism were provided in **Supplementary file S5**.

### 2.9 Functional annotation of *L. plantarum* LPJBC5 based on gene ontology and cluster of orthologues

The genome of LPJBC5 was analyzed for orthologous genes for functional classifications. The functional characterization revealed most of the annotated genes were involved in carbohydrate metabolism (22.28%); amino acid and derivatives (14.58%); protein metabolism (11.93%); co-factor, vitamins, prosthetic group, pigments (8.93%); Nucleotide and Nucleoside metabolism (7.69%); DNA metabolism (5.48%); cell wall and capsules (5.03%); virulence, disease and defense (3.53%).The orthologous proteins were also involved in ion and coenzyme transport, lipid and nucleotide metabolism (**Figure 4**). The complete list of annotated genes and orthologus group, including the COG category and their classification, gene ontology, were provided in the **Supplementary files S6 & S7**.

**Figure 4.**
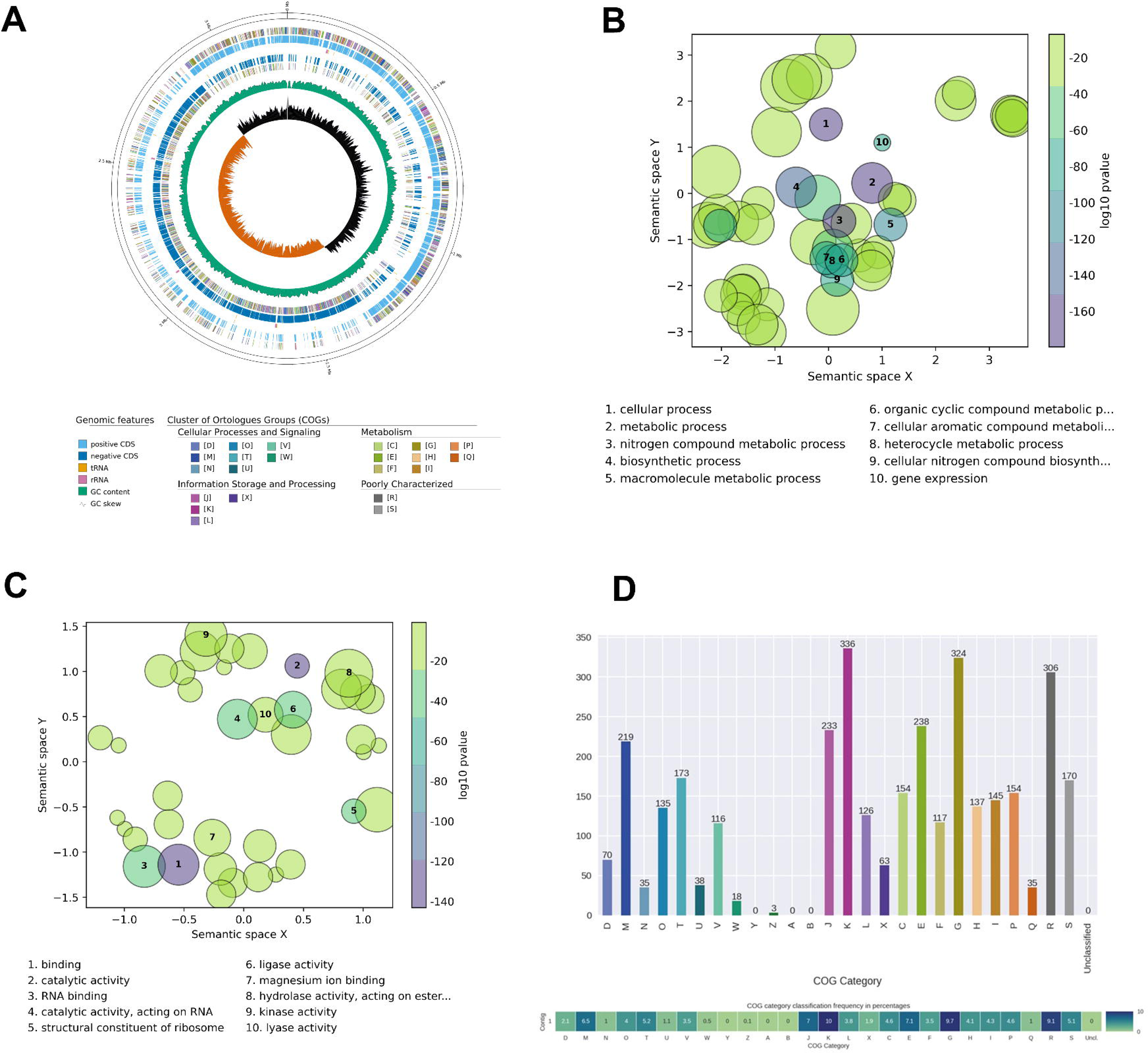
(A) Functional annotation of LPJBC5 genome and its clusters of orthologous groups based on (B) biological process (C) molecular function (D) The frequency of functional orthologous groups detected in the genome of LPJBC5.

## 3. Comparative assessment based on complete genomes of *L. plantarum* strains isolated from various dairy products

We compared the LPJBC5 strain with the existing genomes of *L. plantarum* isolated from dairy products, *viz*. WCSF1 (GCF_000203855.3), ATCC14917(GCF_000143745.1), 12_3 (GCA_004028335.1), DR7 (GCA_003586485.1), YW11(GCA_004028295.1), 13_3 (GCA_004028315.1), TK-P2A(GCA_015377525.1), K25(GCA_003020005.1), LPT52(GCA_023348525.1), ISO1(GCA_026184455.1), IM1130(GCA_925301115.1), IM1131(GCA_925320525.1), IM1155(GCA_925291875.1), C4 (GCA_002994725.1), NL42(GCA_000966475.1), IM930 (GCA_925286555.1), Lp998(GCA_900095055.1), XZ3303(GCA_004123095.1), S2.13(GCA_026156885.1), YW32(GCA_004123035.1), Lp790 (GCA_900095045.1), Lp813(GCA_900095065.1),M92C(GCA_002532175.1), Dad-13(GCA_023547165.1), SKT109(GCA_004025165.1), P-8 (GCA_000392485.2), LZ227(GCA_001660025.1), LZ206(GCA_001659745.1), 10CH (GCA_002005385.2) and DHCU70(GCF_003990985.1). The genome size of all the strains ranges from 3.14 mb to 3.42 mb, with an average GC content of 44%. On average, more than three 3,000 genes were detected across the genome of all strains. The comparative genomic features of these strains are provided in **Supplementary Table S2.**

### 3.1 Pangenomic analysis revealed unique and shared gene clusters of *L. plantarum* strain LPJBC5 and the other related strains

Pangenome analysis revealed that the core genome contained 1844 genes, 254 genes resided in the soft-core region, 1743 genes in the shell region, and 4566 in the cloud region of the genome (**Figure 5A**). A total of 8408 genes were detected during the analysis. In the accumulation curve of *L. plantarum* core and pan-genome, it was observed that the size of the pangenome increases upon the addition of a new genome, whereas the core genome reduces with the addition of a new genome, suggesting an “open” pan-genome nature **(Figure 5B)**. The largest number of accessory genes was 1070, detected in *L. plantarum* IM1155 strain, while the smallest number of accessory genes was 718, detected in *L. plantarum* Dad-13 strain. *L. plantarum* LPJBC5 genome contained 1844 core genes, 893 accessory genes, and 45 species-specific genes (**Figure 5C**). Interestingly, probiotic associated marker genes (approximately 60) were also identified in the core region. Detailed distribution of core, accessory, and unique genes among 31 *L. plantarum* strains was provided in **Supplementary Table S3.**

**Figure 5.**
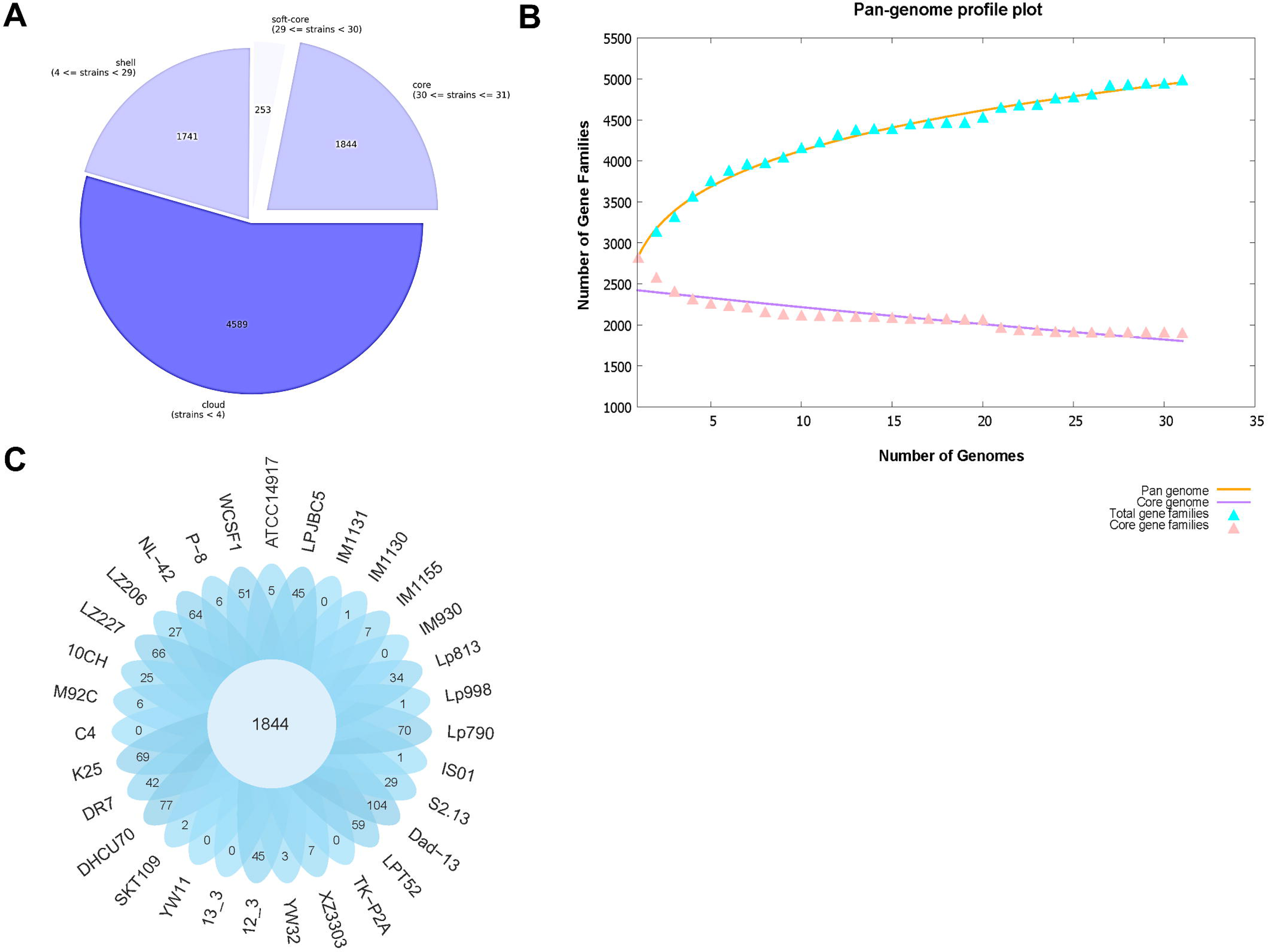
(A) The number of genes belongs to the core, soft core, shell, and cloud region of the *L. plantarum* genome represented. (B) Representation of pangenome profile curves of analyzed *L. plantarum* genomes. The pangenome profile trends were calculated using clustering tools USEARCH. (C) Flower plot diagram showing core and unique genes across all *L. plantarum* strains. The central circle shows the number of genes common to all strains while the petals show the number of genes unique to each strain.

The total gene number of gene families was calculated across all the *L. plantarum* strains. The gene family frequency spectrum presents the number of core, accessory, and unique gene families,1844 core genome families represent 31 genomes, and others are accessory gene families (**Figure 6**).

**Figure 6.**
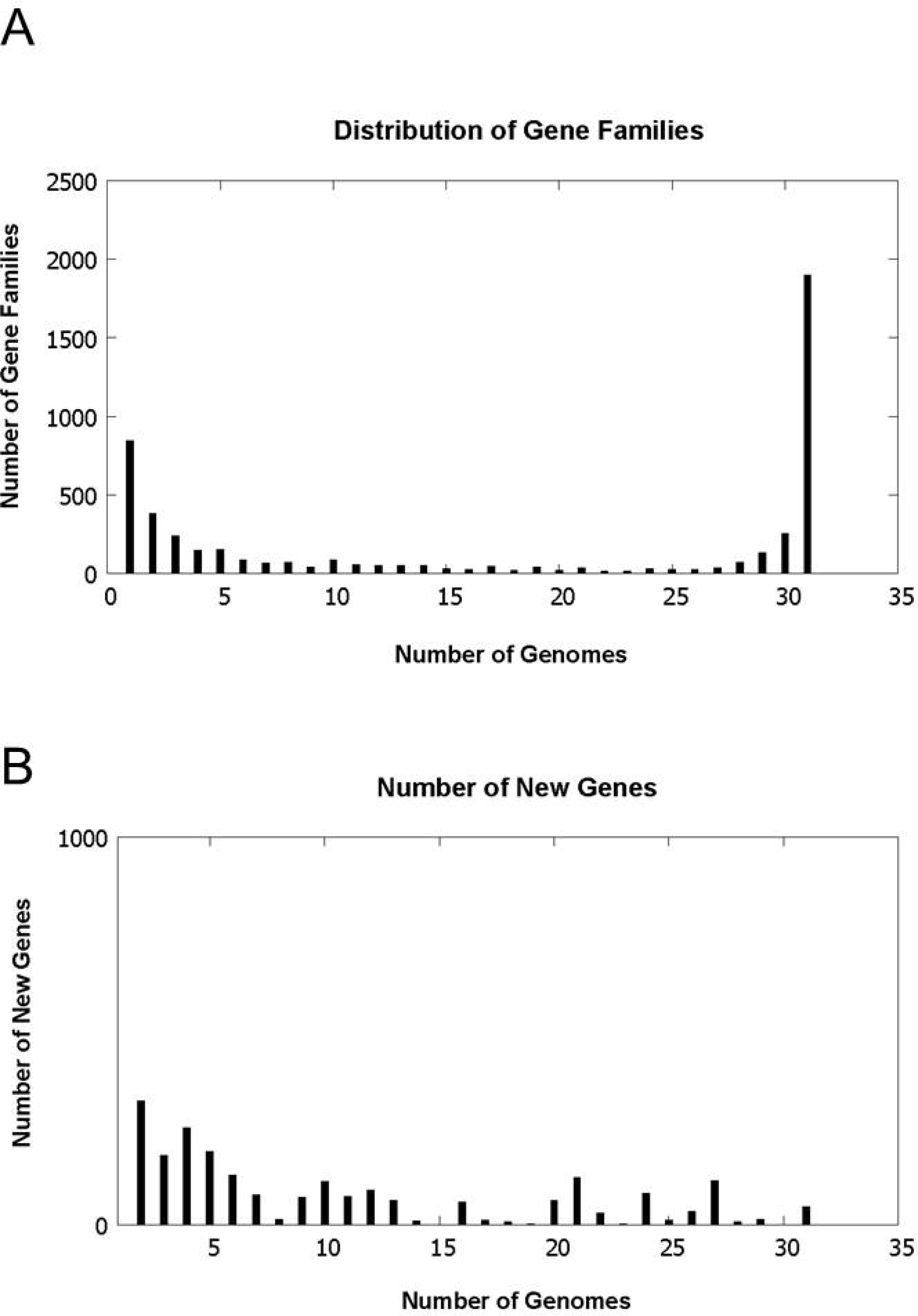
(a) The gene family frequency spectrum of analyzed *L. plantarum* genomes. (b) New gene family distribution after sequential addition of each genome to the analysis.

### 3.2 Phylogenetic analysis of *L. plantarum* genomes

The progressive alignment revealed a synteny among the genomes of all *L. plantarum* strains based on Locally Collinear Blocks (LCBs) (**Supplementary figure S3**). Phylogenetic analysis based on a pan- genomic profile revealed that *L. plantarum* LPJBC5 clustered together with *L. plantarum* [SKT109 &XZ3303] strain. The basal clade consists of ATCC14917 and M92C strains of *L. plantarum*. The clade is further subdivided into multiple clades. 9 strains (K25, YW32, P-8, 13_3, YW11, LZ227, LZ206, S2_13, TK-P2A) from China grouped together with a single strain from Spain (C4), and South Africa (ISO1). 4 strains (IM1130, IM1131, Lp813 and Lp998) from Italy clustered with a single strain from Indonesia (Dad13). A single strain from India (DHCU70) formed a separate clade along with a strain from Ireland (LPT52), United Kingdom (10CH) and Slovenia (WCSF1) (**Figure 7)**.

**Figure 7.**
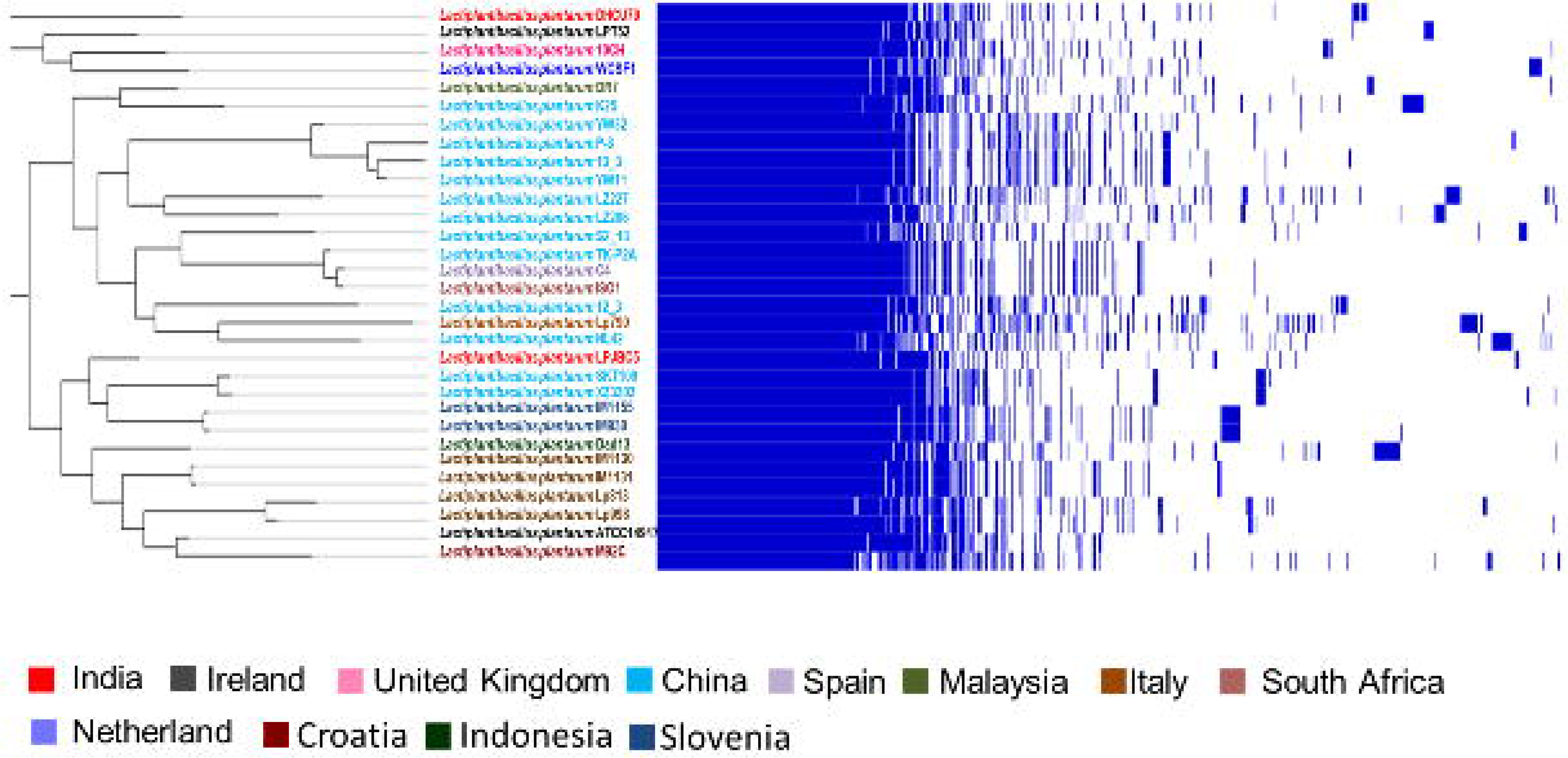
Maximum likelihood-based phylogenetic tree and corresponding pangenome profile based on whole genome sequences of *L. plantarum*. The sequences with a colour code represent the sequences from a particular country.

### 3.3 Functional and pathway mapping of core, accessory, and unique genes across *L. plantarum* LPJBC5 and the other related strains

The distribution of various COG functional categories in core, accessory, and unique gene families was depicted in Figure 11. The result showed that the enriched functions in the core genome of *L. plantarum* strains were associated with metabolism (37%), followed by information storage and processing (30%). Amino acid transport and metabolism (9.68%) and carbohydrate transport and metabolism (8%) were the most enriched metabolic functions. Other enriched metabolic functions include nucleotide transport and metabolism (3.80%), coenzyme transport and metabolism (3.02%), and lipid transport and metabolism (3.13%). Additionally, transcription mechanism (10.85%), translation, ribosomal structure and biogenesis (7.38%), and cell wall/membrane/envelope biogenesis (5.14%) were also enriched in core genes of *L. plantarum* strains. The accessory genes were involved in metabolism (26%), transcription activity (13.97%), cell wall biogenesis (6.60%), cell defense mechanisms (3.33%), replication, recombination, and repair (9.66%). About 14% of the core gene was categorized under general function and unknown function (8.95%). Similarly, for accessories and unique genes, around 24% of the total genes were grouped under general function and poorly characterized (12%) (**Figure 8**).

**Figure 8.**
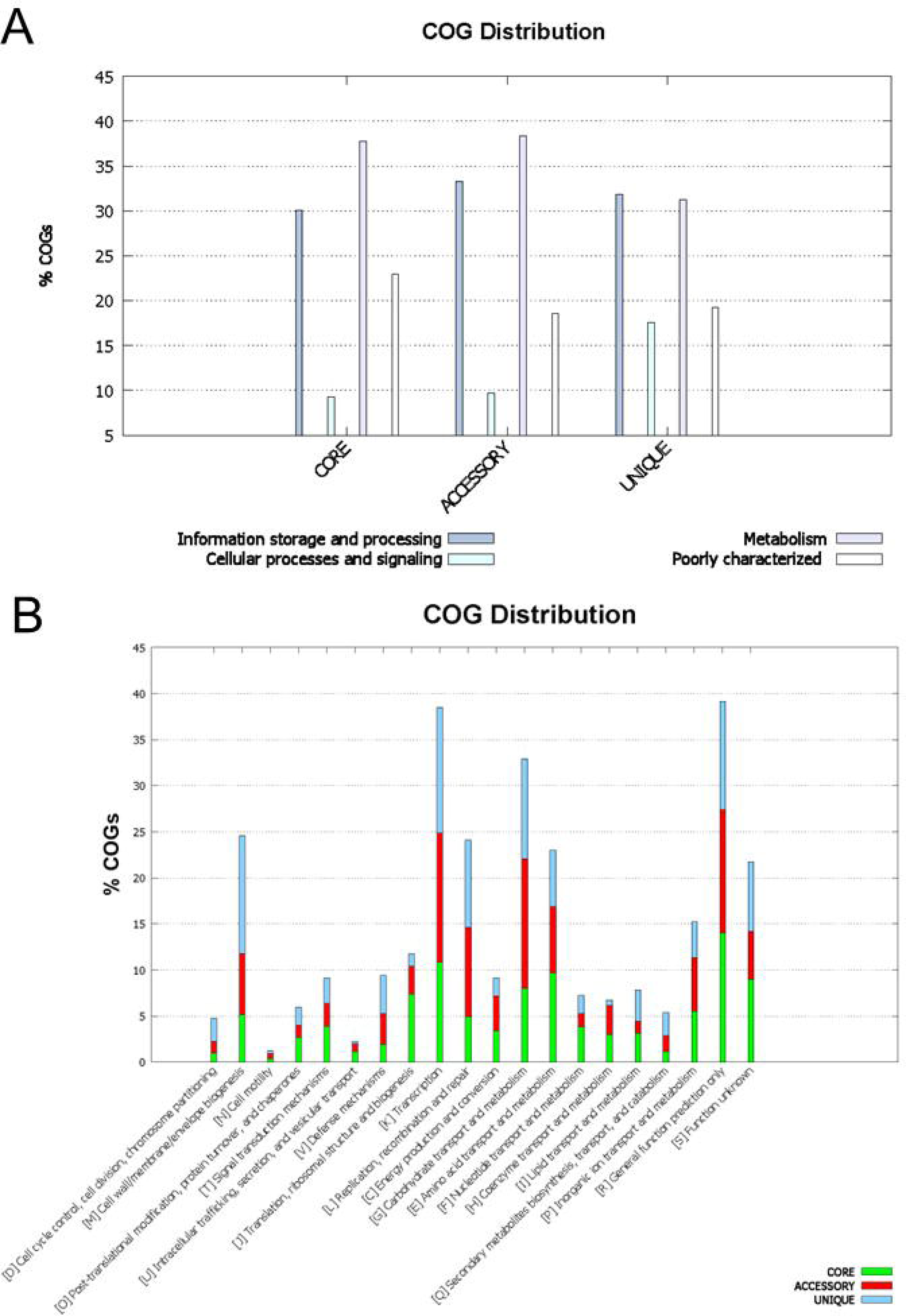
(A) Overall COG distribution of core, accessory, and unique genes. (B) Detail distribution of COG category of core, accessory, and unique genes of *L. plantarum* pan genomes.

KEGG pathway mapping revealed overall dominant representation of metabolism related pathways (**Figure 9A**) which is also supported by COG distribution regarding metabolic function. In the sub-category of metabolism, the most abundant function in the core and accessory genome conserved carbohydrate metabolism (**Figure 9B**). The details of mapped pathways of core and accessory and unique genes across *L. plantarum* genomes were provided in a **supplementary file S8.**

**Figure 9.**
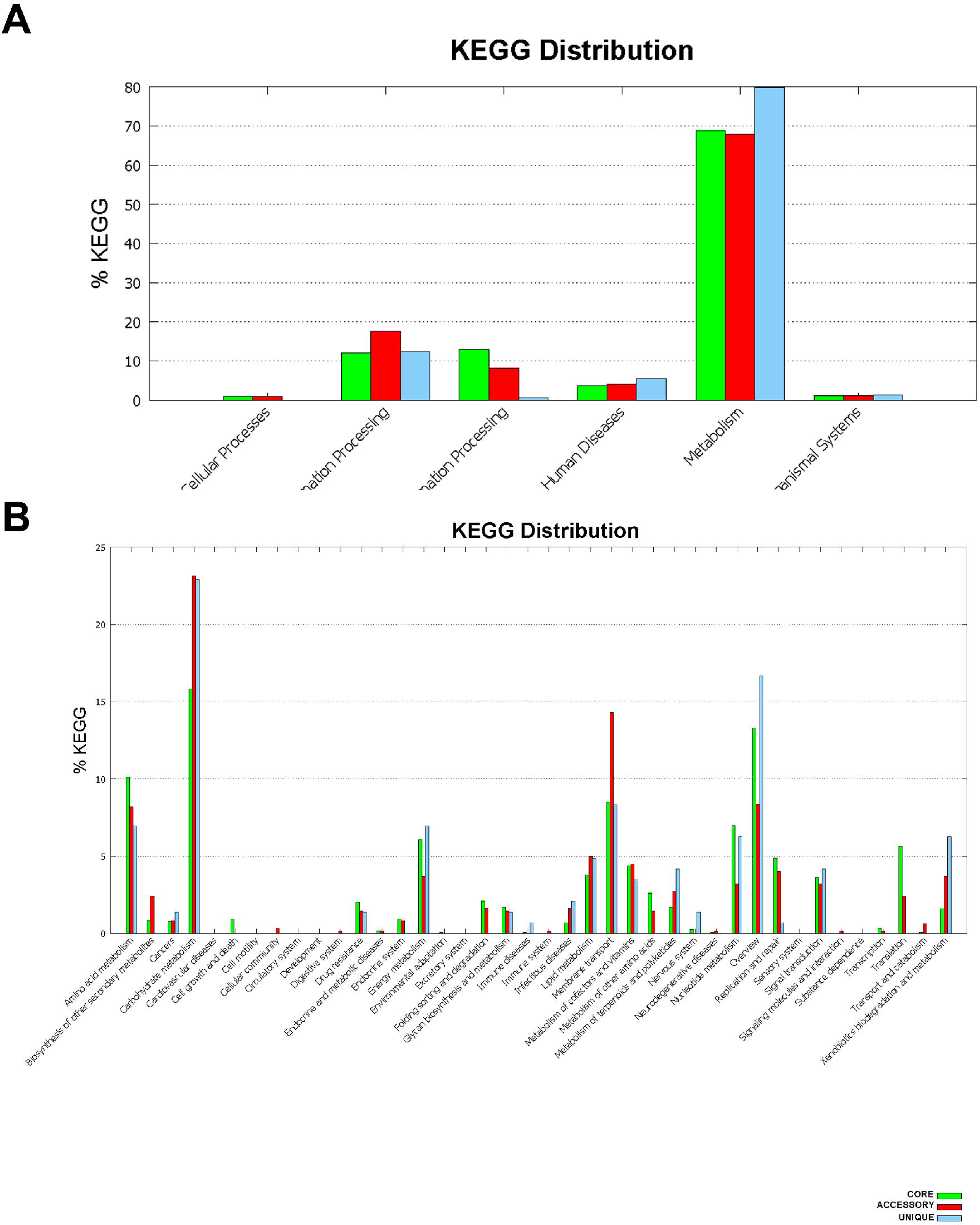
(A) Overall KEGG pathway distribution of core, accessory, and unique genes. (B) Detail distribution of KEGG pathways of core, accessory, and unique genes of *L. plantarum* pan genomes. (C) Significantly enriched pathways (Top 20) of the *L. plantarum* genomes.

### 3.4 **Identification of mobile genetic element (MGEs)**

Microorganisms emerge through DNA mutations and the transfer of mobile genetic elements contributing to their adaptation and evolution. MGEs often transfer various accessory genes that give their host cell advantages including antibiotic resistance, horizontal gene transfer and virulence factors. In this study, a total of 20 MGEs were predicted, of which 17 were insertion sequences, and 3 were composite transposons. The insertion sequences and transposons belong to ISP1 and ISP2 groups in *L. plantarum* LPJBC5. During comparative analysis, the maximum number of MGEs was detected in *L. plantarum* YW11 and *L. plantarum* 13_3 strains, while no MGEs were detected in *L. plantarum* M92C strain (**Figure 10).** The complete list of detected MGEs along with their contig family and position in the genome were provided in **supplementary file S9**. Furthermore, we identified copy number of genes across 31 *L. plantarum* genomes. As shown in Fig. 12b, we compared the distribution of genes among the *L. plantarum* strains. The number of single-copy genes was 1002, while 548 multiple-copy genes were detected in the genome of LPJBC5.

**Figure 10.**
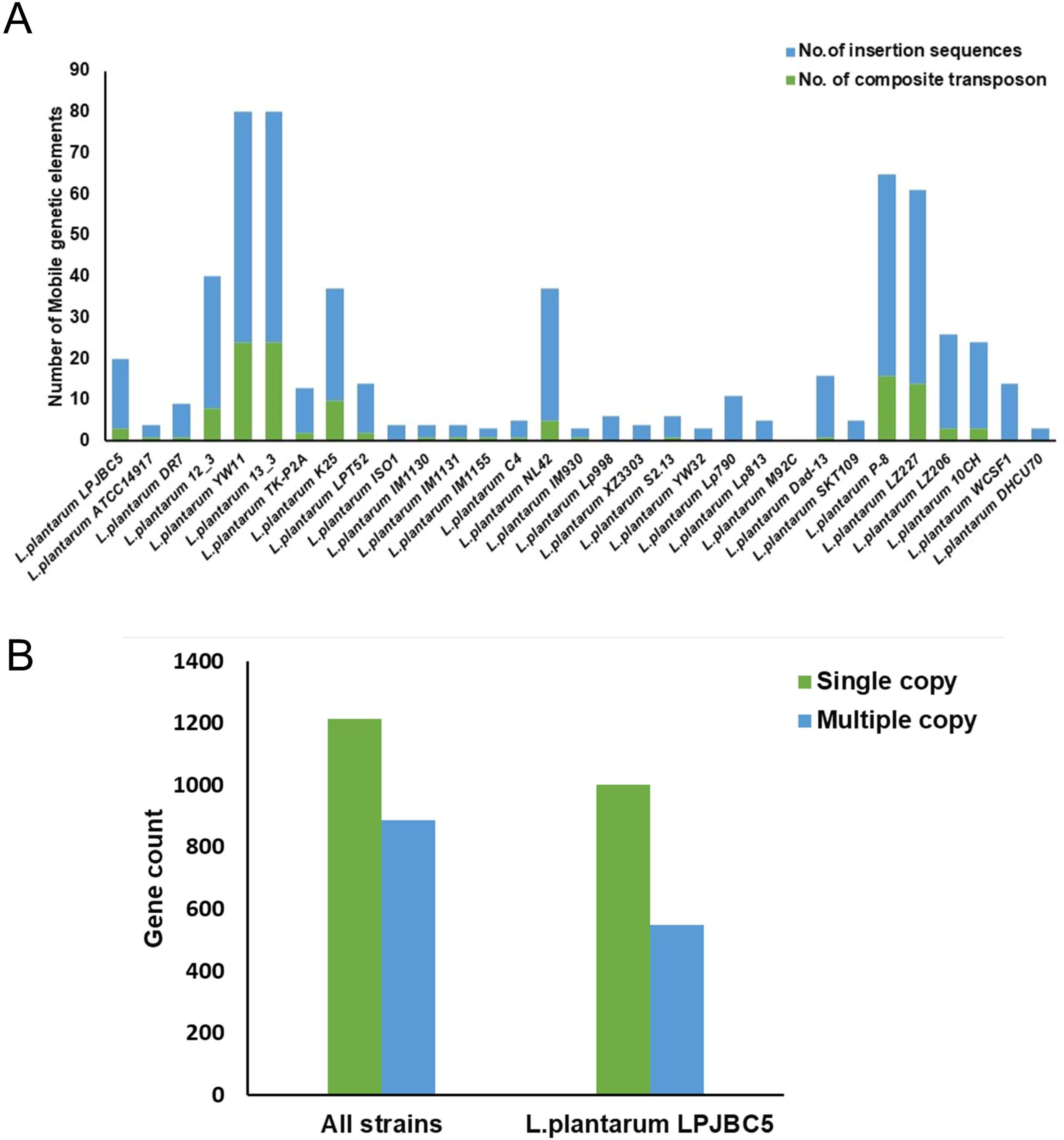
(A) The stacked bar plot represented predicted mobile genetic elements (MGEs) for L. plantarum strains’ genomes. Blue highlighted the insertion sequences, and green highlighted the composite transposons. (B) Distribution of single and multiple copies of genes in *L. plantarum* LPJBC5 and other strains.

## Discussion

*Lactiplantibacillus plantarum* (formerly known as *Lactobacillus plantarum*) is a well-documented probiotic lactic acid bacterium with a wide range of applications in the food and health sectors (Corsetti and Valmorri et al. 2011, Fidanza, Panigrahi et al. 2021). With the advent of whole-genome sequencing, our understanding of the genetic basis of its probiotic attributes has expanded remarkably. *Lactiplantibacillus plantarum* LPJBC5 was previously reported to harbor probiotic and anti-aging properties (Kumar et al. 2022). However, the genes responsible for their characteristics were not reported. Therefore, this study has meticulously focused on the complete genome of LPJBC5 and found that significant genes were involved in carbohydrate metabolism, synthesis of exopolysaccharides, adhesion, genes responsible for immunomodulatory properties, bacteriocin productions, antibiotic resistance, reducing oxidative stress, acid/bile resistant and longevity.

Whole genome analysis revealed that LPJBC5 has a strong metabolism for carbohydrates and amino acids. Genes are involved in the metabolism of starch, glycogen, cellobiose, alpha-mannan, glycogen, galactan, and beta-glucoside. The genome of LPJBC5 had dominance of genomic GH, which suggests that the strain can efficiently utilize different carbon sources, which further pave the way for the formation of nucleotide sugar bases. Additionally, GT family is one of crucial gene family for EPS biosynthesis (Zeidan et al 2017). Glycosyl transferase catalyze the transfer of sugars from the activated donor molecules to specific acceptors and are essential for the formation of surface structures recognized by host immune system (Venter, Meyer et al. 2022). Previous study reported that the GT2 and GT4 enzymes are involved in the biosynthesis of sugars such as sucrose, cellulose, lipopolysaccharide and chitosan which are further associated with the water holding capacity and viscosity of the final fermented milk product (Oehme et al. 2019). The above results suggests that LPJBC5 has efficient sugar biosynthesis properties. Previously Zhao & Liang et al (2022) has reported that MC5 having similar sugar biosynthesis properties improved rheological properties of yogurt.

Exopolysaccharides (EPS) produced by lactic acid bacteria, such as *L. plantarum*, have gained significant attention in recent years due to their potential health benefits and applications in the food industry(Silva, Lopes Neto et al. 2019). The genomic analysis revealed the presence of EPS-producing genes (*eps* gene cluster, Mur ligase gene family) in the genome of LPJBC5. In addition, strain possesses a variable region that specifically includes oligosaccharide flippase family protein (*wzx*) and polysaccharide polymerase (*wzy*). The genes *eps*C, *eps*D, *eps*F, *eps*H, and *eps*B in LPJBC5 play crucial roles in the biosynthesis and export of EPS. The protein encoded by *eps*C helps in transferring the initial sugar residue and helps in initiation of EPS synthesis. The *eps*D gene encodes for a polymerase enzyme that facilitates the elongation of the EPS chain by adding sugar units sequentially, forming a long carbohydrate chain. Further, the *eps*F enzyme is involved in the secretion and ensures EPS transportation outside the cells. The EPS structure is modified by *eps*H gene by adding specific side chains to the primary EPS structure, thus imparting specific functional attributes. The *eps*B enzyme is crucial in regulating EPS production and response to environmental cues. Previous reports suggests that EPS has huge implication in promoting gut health and are also an important prebiotic (Zeidan, Poulsen et al. 2017, Zhao, Liang et al. 2023).

The LPJBC5 genome harbor numerous adhesion-related genes, like Mucin binding domain (MucBP) and mucus adhesion promoting protein genes (*mapA*), that potentially allowing the bacterium to adhere to the intestinal mucosa, enhancing its colonization . The genome of LPJBC5, possesses fibronectin binding proteins (fbp), sortase (*srt*A), enolases, LPXTG anchored proteins, *exo*A, and *lsp*A. Earlier findings indicated that the MucBP domains within the Mub protein initiate adhesion to the mucin on the surface of epithelial cells. They play a crucial role in the adherence of microbial species to the mucosal surfaces in host organisms(MacKenzie, Jeffers et al. 2010). These proteins specifically recognize and bind to mucins, which are heavily glycosylated proteins found in mucus layers covering epithelial surfaces, such as those in the respiratory, gastrointestinal, and urogenital tracts(McGuckin, Lindén et al. 2011). In addition to their role in adhesion, mucin-binding proteins can influence other cellular processes and contribute to cell signaling and communication at the cell surface(Singh and Hollingsworth et al. 2006).

Genome analysis revealed the presence of specific genes (*dlt*B, *dlt*D) that might play a role in modulating host immune responses(Lebeer, Vanderleyden et al. 2008). The *dlt* operon comprises five genes: *dlt*X, *dlt*A, *dlt*B, *dlt*C, and *dlt*D. In *L. plantarum* mutants, removing the *dlt*B and *dlt*D genes resulted in elevated production of the anti-inflammatory cytokine IL-10 compared to the wild type strain in co-culture experiments(Grangette, Nutten et al. 2005).

Cellular metabolism can lead to an overproduction of reactive oxygen species (ROS) and reactive nitrogen species (RNS), inducing oxidative stress and subsequently damaging macromolecules (Lu and Holmgren et al. 2014). The genome of *L. plantarum* LPJBC5 features genes, such as *clp*C, *clp*E, *clp*B, and *clp*X, that respond to the stress and safeguard proteins from undue damage. Recognizing probiotics’ remarkable potential as antioxidants and their role in preserving intestinal redox balance, our research focused on antioxidant-associated genes in the genome. Key genes from the thioredoxin system, including *trx*A and *trx*B, were identified. This system supplies electrons to thiol-dependent peroxidases, aiding in rapidly removing ROS and RNS(Lu and Holmgren et al. 2014). *L. plantarum* LPJBC5’s genome exhibits a multitude of genes suggesting a natural evolution to withstand and adapt to acidic conditions and bile challenges. Specific genes, like *cbh*, *opp*A, and *dps*, are pivotal in this adaptive tolerance. Furthermore, the genome encodes NADH oxidation-associated genes, such as the nox gene (NADH oxidase) and *npr* genes (NADH peroxidase). Both catalase and NADH oxidase/peroxidase are integral for neutralizing hydrogen peroxide and other ROS. The genome also possesses the gene for glutathione reductase, a vital antioxidant enzyme that upholds glutathione levels, a primary antioxidant compound. The thioredoxin system provides electrons to thiol-dependent peroxidases to counteract ROS. Additionally, the *asp*B gene in the *L. plantarum* LPJBC5 genome encodes Aspartate aminotransferase. This enzyme’s role in transforming oxoglutarate into glutamate further augments the genome’s resistance to oxidative stress.

Secondary metabolites are small organic molecules with diverse biological functions (Medema, Blin et al. 2011, Abegaz and Kinfe et al. 2019). This study explores the potential of the probiotic strain L. plantarum LPJBC5 to produce putative metabolites. Four secondary metabolites biosynthetic gene clusters were identified, including terpene, T3PKS, Ribosomally synthesized and post-translationally modified peptide product (Ripp-Like), and cyclic-lactone-autoinducer. The genome of *L.plantarum* LPJBC5 contained T3PKS with the *mva*S gene, encoding hydroxymethylglutaryl-CoA synthase. This enzyme is sensitive to feedback substrate inhibition by acetoacetyl-CoA, built up from pyruvate in the colluding steps of glycolysis. The genome also contains additional biosynthetic genes encoding putative HAD hydrolase, D-lactate dehydrogenase involved in pyruvate metabolism. These are involved in NAD+ regeneration during lactic acid fermentation (Goffin, Lorquet et al. 2004, Kasai, Suzuki et al. 2019). It was reported that they are also involved in adhesion, absorption, and believed to have safe additives for application in food fermentation(Jatuponwiphat, Namrak et al. 2019, Soumya and Nampoothiri et al. 2021). The genome contains Ripp-like biosynthetic gene cluster with *lag*D gene. The gene encodes Lactococcin G-processing and transport ATP binding protein, a bacteriocin capable of exhibiting antimicrobial activity (Ishibashi, Zendo et al. 2015). Because of their heat stability, and pH tolerance, bacteriocins from LAB might be used as preservatives to prevent spoilage and growth of pathogens in food (Ibrahim, Ayivi et al. 2021). *L. plantarum* strains isolated from different environments have been reported to produce both lantibiotic and/or non-lantibiotic bacteriocins(González, Arca et al. 1994, Rekhif, Atrih et al. 1995, Franz, Toit et al. 1998, Leal- Sánchez, Jiménez-Díaz et al. 2002, Field, Hill et al. 2010, Domínguez-Manzano and Jiménez-Díaz et al. 2013, Song, Zhu et al. 2014, Todorov, Holzapfel et al. 2016). The genome of *L. plantarum* LPJBC5 contains genes from the Plantaricin family, such as PlantaricinK (*pln*K), PlantaricinJ (*pln*J), PlantaricinN (*pln*N), PlantaricinA (*pln*A), PlantaricinF (*pln*F), and PlantaricinE (*pln*E). Moreover, the genes *pln*O, *pln*C, *pln*D, *pln*S, and *pln*L are also present. Bacteriocin production is activated through the phosphorylation of *pln*C/*pln*D when the external concentration of *pln*A reaches a certain threshold (Diep, Johnsborg et al. 2001). The genome features immunity-related proteins like *pln*L, *pln*E, *pln*F, and *pln*S, and ATP-binding and permease proteins such as *pln*G and *pln*O. Earlier research has highlighted the importance of bacteriocins in preserving food, as they counteract the growth of microbes responsible for food spoilage and pathogenicity (Silva, Silva et al. 2018, Ng, Zarin et al. 2020).

A comprehensive pangenome analysis was performed to discern the genomic differences, identify both core and accessory genes and their functional elucidation within *L. plantarum* strains isolated from fermented dairy items retrieved from repositories. Few studies focused on the pangenome of *L. plantarum*(Carpi, Coman et al. 2022). It was observed that a total of 86 probiotic marker genes were identified of which 76 genes belong to core/soft-core regions, 8 genes in shell region and 2 in cloud region. Previous reports suggests that 70 probiotic marker genes associated with stress resistance, adhesion, active metabolism and gut persistence were identified in core/soft-core region. In comparison, 5 probiotic marker genes (*bsh*A, *opp*A, *srt*A, *xyl*A, *gla*_2) belong to shell and cloud region (Carpi, Coman et al. 2022). 893 accessory genes were identified in LPJBC5 while 911 and 943 accessory genes were reported in *L. plantarum* ATCC 14917 and *L. plantarum* WCSF1 (Carpi, Coman et al. 2022) respectively.

The phylogenetic relationship revealed that LPJBC5 clustered together with the genome of *L. plantarum* (SKT09 and XZ3303) while clustered distinctly with *L. plantarum* DHCU70. *L. plantarum* SKT09 was isolated from kefir grain and was reported to inhibit aging in mice. Interestingly, Kumar (2022) has previously reported the anti-aging property of LPJBC5 in *C. elegans*. The genomic analysis also supports longevity-associated genes in LPJBC5, including *kat*E, CAT, *cat*B and *srp*A. Aging is a complex process of accumulation of molecular, cellular, and organ damage, leading to loss of function and increased vulnerability to disease and death (Kumar, Joishy et al. 2022). *L. plantarum* DHCU70 was isolated from naturally fermented milk product (Rai, Shangpliang et al. 2016) while LPJBC5 was isolated from boiled milk curd (Joishy, Dehingia et al. 2019, Kumar, Joishy et al. 2022) which give insights into the importance of source of probiotic bacteria isolation.

During analysis, a total number of 846 strain specific genes were detected across the *L. plantarum* genomes. High number of strain specific genes were observed in *L. plantarum* Dad-13 (104), *L. plantarum* Lp790 (70), *L. plantarum* K25 (69); while very less strain specific genes were observed in *L. plantarum* ATCC14917 (5), *L. plantarum* P-8 (6), *L. plantarum* M92C (6), *L. plantarum* SKT109 (2), *L. plantarum* YW32 (3), *L. plantarum* XZ3303 (7), *L. plantarum* IM1155 (7), *L. plantarum* IS01 (1), *L. plantarum* Lp998 (1), *L. plantarum* IM1130 (1). However, no unique genes were detected in *L. plantarum* C4, *L. plantarum* YW11, *L. plantarum* 13_3, *L. plantarum* TK-P2A, *L. plantarum* IM930, *L. plantarum* IM1131. 45 unique genes were detected in the *L. plantarum* LPJBC5 genome.

The COG-based enrichment analysis showed that the core genomes among 31 *L. plantarum* strains were primarily enriched with metabolism, information storage, and processing. Previously, the metabolism related genes were also reported to be present in the core region (Carpi, Coman et al. 2022). The accessory genomes were enriched with metabolism, transcription activity, and cell wall biogenesis. In *the L. plantarum LPJBC5 genome, the genes were majorly involved in carbohydrate metabolism and transport, transcription, translation, post-translational modification, protein turnover, chaperone functions,* and cell wall/membrane/envelop biogenesis. A higher number of COGs related to carbohydrate metabolism could indicate the bacterium’s adaptability in utilizing different sugars, aligning with its role in diverse fermentative processes. Comparative studies of L. plantarum strains from fermented dairy products showed a substantial gene overlap. Pangenome analysis highlighted significant genomic variability within *L. plantarum*, suggesting various genetic content across strains. The core genome evaluation identified a consistent set of genes in all strains, probably underpinning essential cellular activities and metabolic roles.

Mobile genetic elements are critical parameters in defining a strain’s probiotic potential because they contribute to antibiotic resistance and horizontal gene transfer (Carpi, Coman et al. 2022). Additionally, they play a pivotal role in the evolution and adaptation, for thriving in a ecological niche (Evanovich, de Souza Mendonça Mattos et al. 2019). In *L. plantarum* LPJBC5 genome, mobile genetic elements, insertion sequence, and transposon were detected. While compared with other strains of *L. plantarum*, we detected a range of transposable elements and insertion sequences. The MGE depicted the presence of antibiotic resistance genes against Vancomycin showing 31.93% of identity. *L. plantarum* JBC5 comply with the safety and efficacy requirements for probiotic properties provided by Indian council of medical research (ICMR), European Food Safety and Standards (EFSA) and U.S. Food and Drug Administration. LPJBC5 belong to generally regarded as safe group, their genomes are fully sequenced and found significant genes associated with the probiotic properties and longevity. The strain has no pathogenicity which was confirmed using Pathogen finder tool. These suggests that LPJBC5 can be used as a starter in food industries.

## Conclusion

*Lactiplantibacillus plantarum* LPJBC5 showcases potential as a probiotic with antioxidative properties and exopolysaccharide synthesis capabilities. This study delves into the genomic landscape of *L. plantarum* LPJBC5, contrasting its genetic attributes with 30 *L. plantarum* strains isolated from dairy products. By the full genome sequence and distinct genomic characteristics, we hope to deepen the comprehension of LPJBC5’s probiotic actions and potential uses. However, further validation through *in vivo* and *in vitro* analyses remains crucial.

## Data Availability

The whole genome data were uploaded in NCBI SRA against the Bioproject ID PRJNA1085560.

## Funding

This research was funded by the ST/SC community development programme in IASST (SEED/TITE/2019/103), Department of Science and Technology (DST), Govt. of India.

## Author contribution

Anupam Bhattacharya: Data curation, Formal analysis, Investigation, Methodology, Visualization, Writing – original draft, Writing – review and editing ; Tulsi Joishy: Investigation, Methodology, Writing – review and editing; Mojibur R. Khan: Conceptualization, Funding acquisition, Resources, Supervision, Writing – review and editing.

## Conflicts of Interest

The authors declare no conflict of interest.

## Supplementary Materials

**Supplementary file S1.** Dataset description of publicly available *L. plantarum* genome.

**Supplementary file S2.** Identification of bacterial strain based on Ribosomal multilocus sequence typing(rMLST).

**Supplementary file S3.** Detected Carbohydrate-Active EnZymes (CAZymes) in the genome of *L. plantarum* LPJBC5.

**Supplementary file S4.** A list of putative bacteriocin operons is presented in the genome.

**Supplementary file S5. A** list of antimicrobial resistance genes detected in the genome.

**Supplementary file S6.** Annotation of *L. plantarum* LPJBC5 genome.

**Supplementary file S7.** Significantly enriched biological processes and molecular functions based on gene ontology.

**Supplementary file S8.** Mapped pathways of core, accessory, and unique genes across *L. plantarum* genomes.

**Supplementary file S9.** List of detected MGEs along with contig family and position in the genome of all *L. plantarum* genomes.

**Supplementary figure S1.** Longevity regulating pathways.

**Supplementary figure S2.** Detection of putative secondary metabolites in *L. plantarum* LPJBC5 genome.

**Supplementary figure S3.** The progressive alignment of all genomes of *L. plantarum* strains.

**Supplementary Table S1.** Average nucleotide identity (ANI).

**Supplementary Table S2.** Comparative genome summary of *L. plantarum* strains.

**Supplementary Table S3.** Distribution of core, accessory, and unique genes among 31 *L. plantarum* strains.

## Notes

### Competing Interest Statement

The authors have declared no competing interest.

