## Supplementary figure S1 for "Exploring the probiotic potential, antioxidant capacity, and healthy aging based on whole genome analysis of *Lactiplantibacillus plantarum* LPJBC5 isolated from fermented milk product"

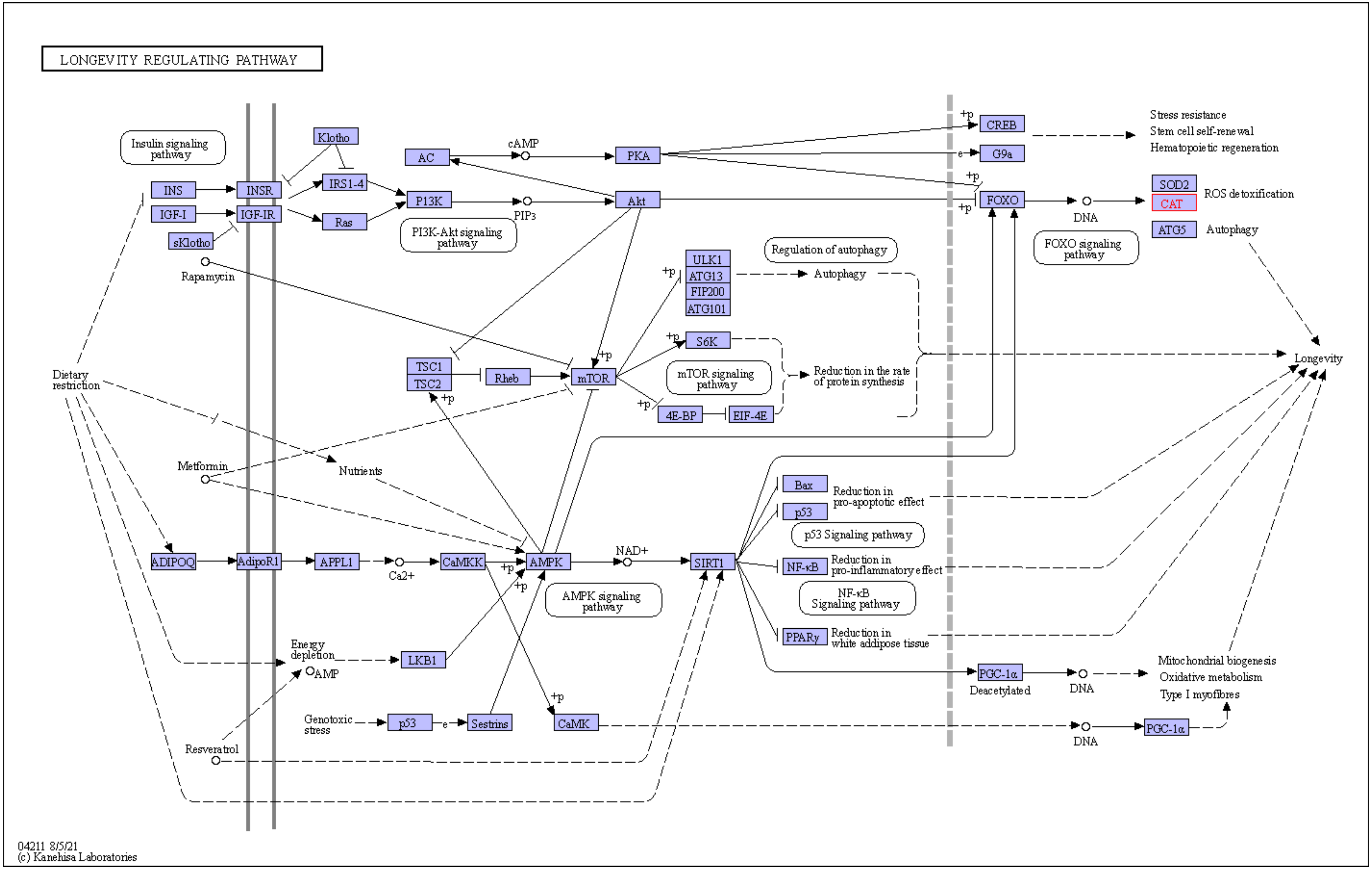


**Supplementary figure S1.** Involvement of *kat*E gene in longevity regulating pathway(map04211)

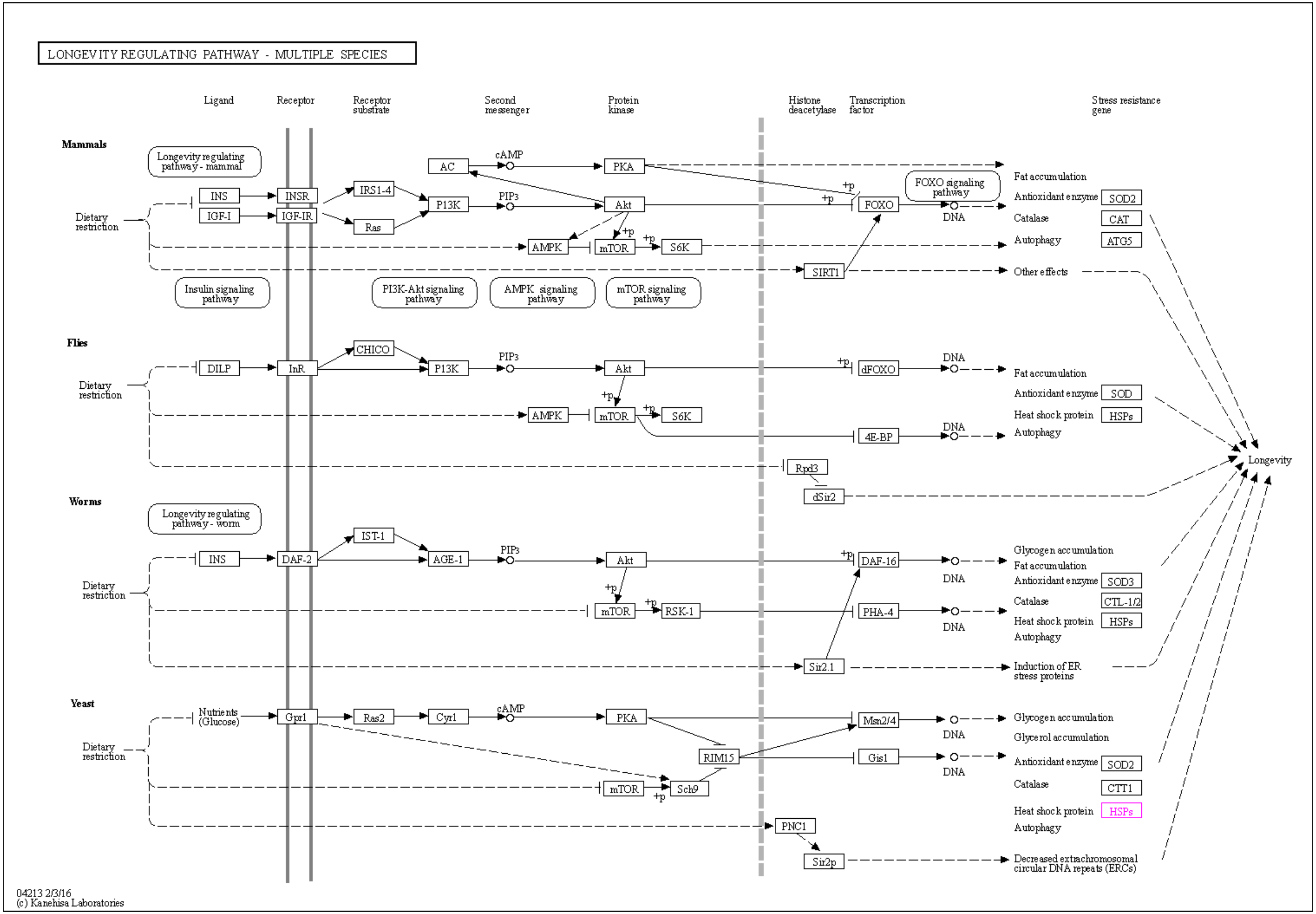


**Supplementary figure.** Association of *clp*B (HSP) gene in longevity regulating pathway (map04213).

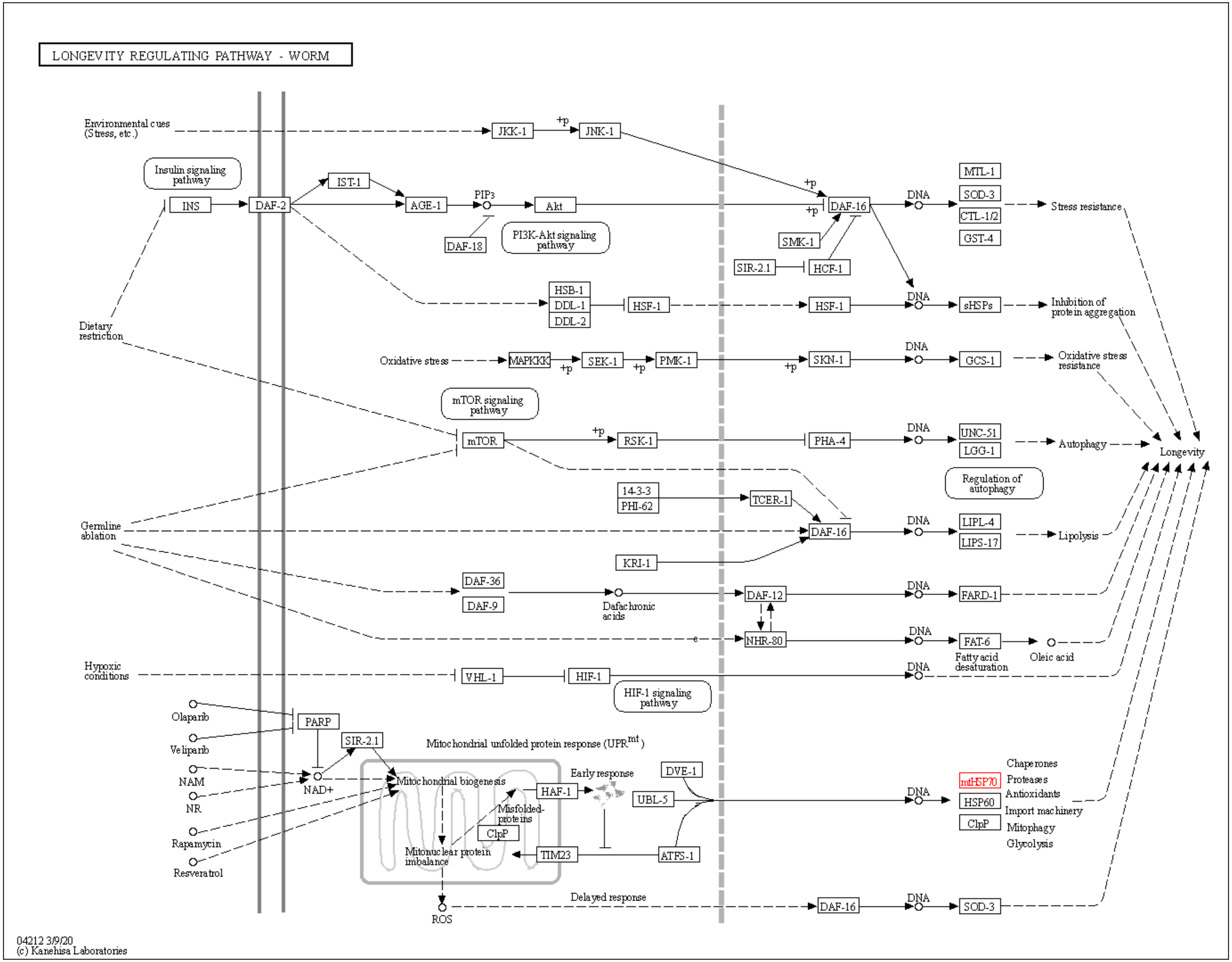


**Supplementary figure S1.** Association of *dna*K(HSP70), *gro*EL (HSP60) gene in longevity regulating pathway (map04043).
