## Supplementary figure S2 for "Exploring the probiotic potential, antioxidant capacity, and healthy aging based on whole genome analysis of *Lactiplantibacillus plantarum* LPJBC5 isolated from fermented milk product"

| Region | Type | From | To |
|---|---|---|---|
| Region 1 | RiPP-like | 19,991 | 32,141 |
| Region 2 | T3PKS | 1,483,836 | 1,525,005 |
| Region 3 | terpene | 2,487,366 | 2,508,247 |
| Region 4 | cyclic-lactone-autoinducer | 2,757,616 | 2,778,321 |

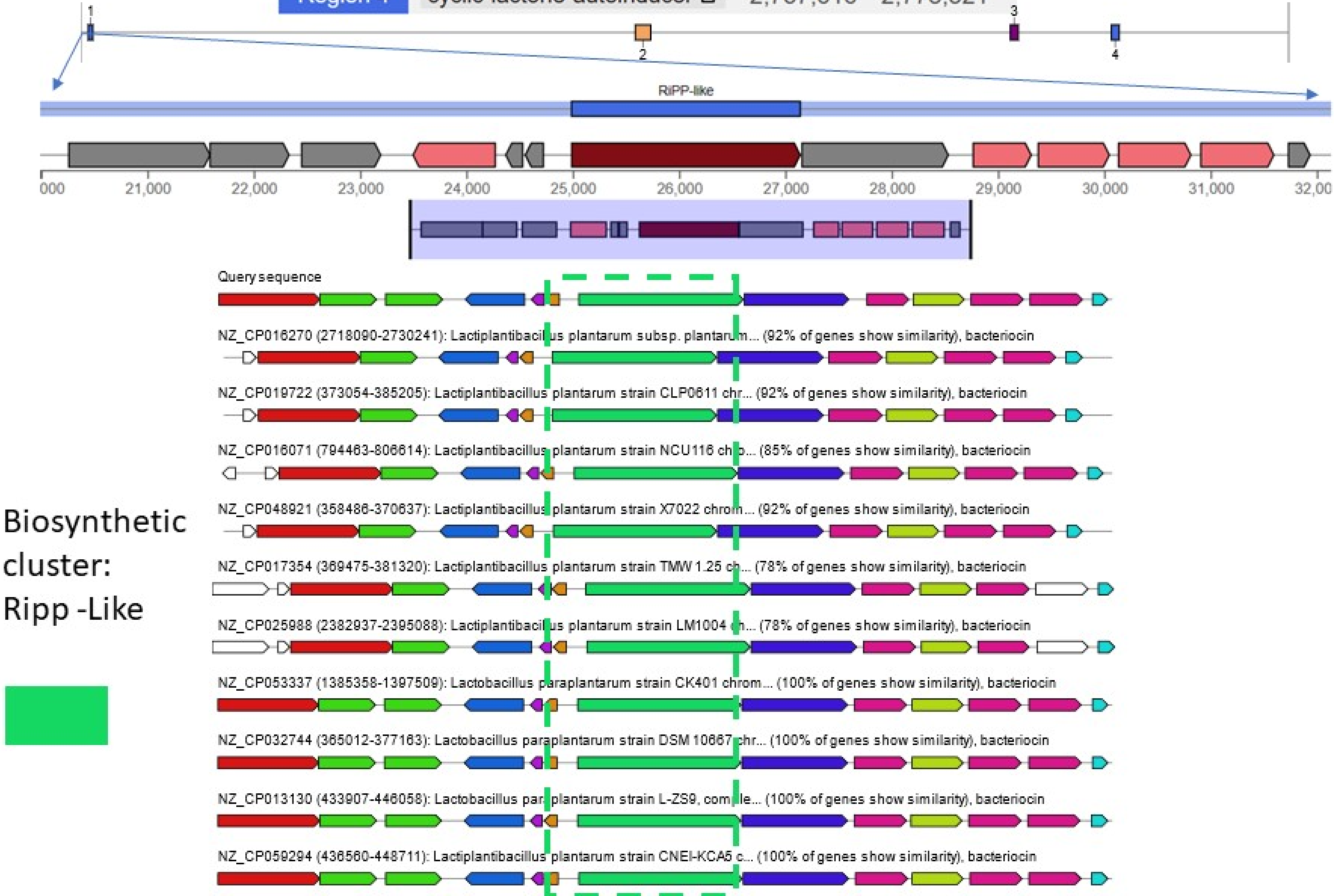

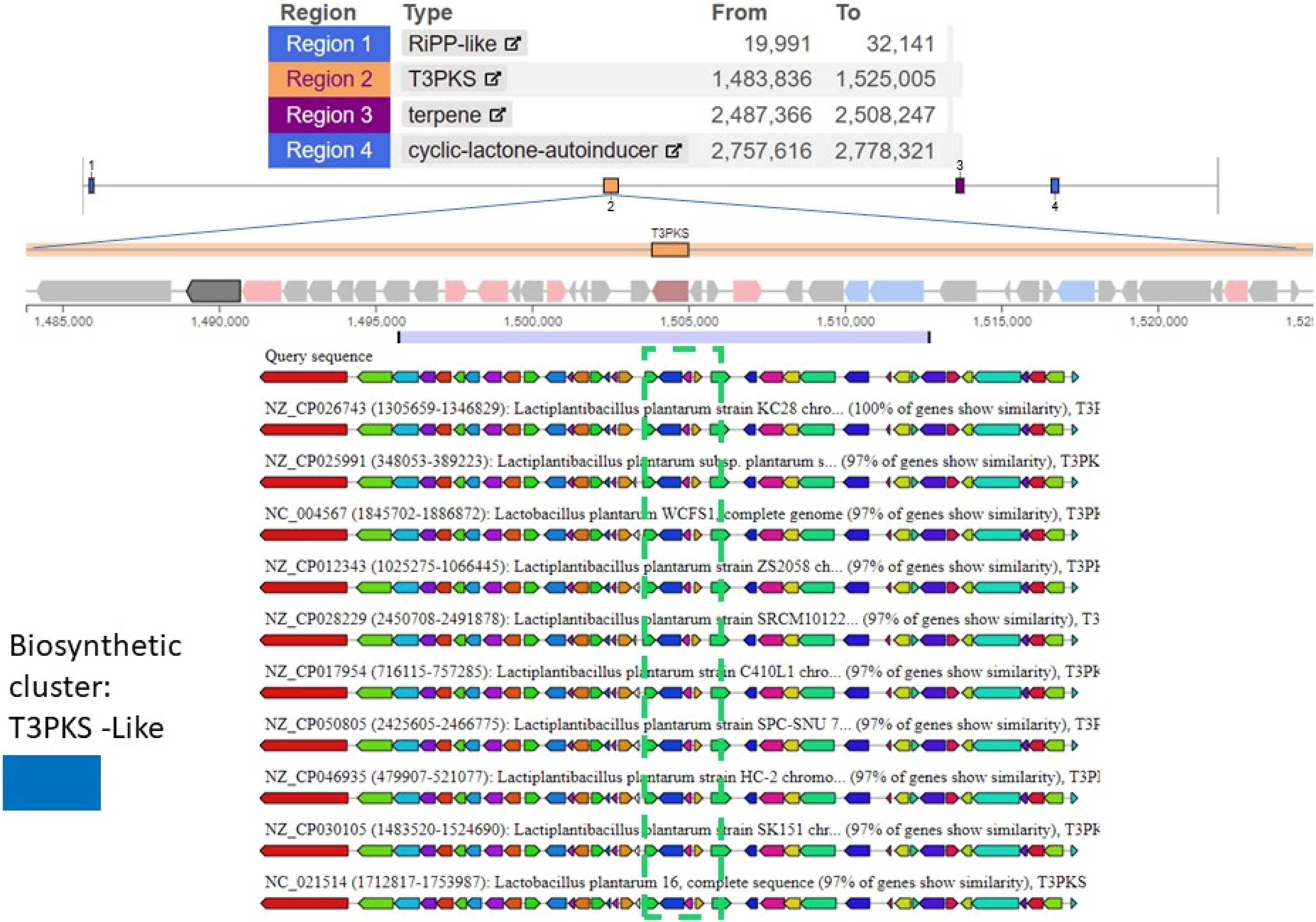
Region | Type | From | To
Region 1 | RiPP-like | 19,991 | 32,141
Region 2 | T3PKS | 1,483,836 | 1,525,005
Region 3 | terpene | 2,487,366 | 2,508,247
Region 4 | cyclic-lactone-autoinducer | 2,757,616 | 2,778,321
T3PKS
1,485,000
1,490,000
1,495,000
1,500,000
1,505,000
1,510,000
1,515,000
1,520,000
Query sequence
NZ_CP026743 (1305659-1346829): Lactiplantibacillus plantarum strain KC28 chro... (100% of genes show similarity), T3I
NZ_CP025991 (348053-389223): Lactiplantibacillus plantarum subsp. plantarum s... (97% of genes show similarity), T3PK
NC_004567 (1845702-1886872): Lactobacillus plantarum WCFS1, complete genome (97% of genes show similarity), T3PI
NZ_CP012343 (1025275-1066445): Lactiplantibacillus plantarum strain ZS2058 ch... (97% of genes show similarity), T3PI
NZ_CP028229 (2450708-2491878): Lactiplantibacillus plantarum strain SRCM10122... (97% of genes show similarity), T3
NZ_CP017954 (716115-757285): Lactiplantibacillus plantarum strain C410L1 chro... (97% of genes show similarity), T3PK
NZ_CP050805 (2425605-2466775): Lactiplantibacillus plantarum strain SPC-SNU 7... (97% of genes show similarity), T3I
NZ_CP046935 (479907-521077): Lactiplantibacillus plantarum strain HC-2 chromo... (97% of genes show similarity), T3PI
NZ_CP030105 (1483520-1524690): Lactiplantibacillus plantarum strain SK151 chr... (97% of genes show similarity), T3PK
NC_021514 (1712817-1753987): Lactobacillus plantarum 16, complete sequence (97% of genes show similarity), T3PKS
Biosynthetic
cluster:
T3PKS -Like

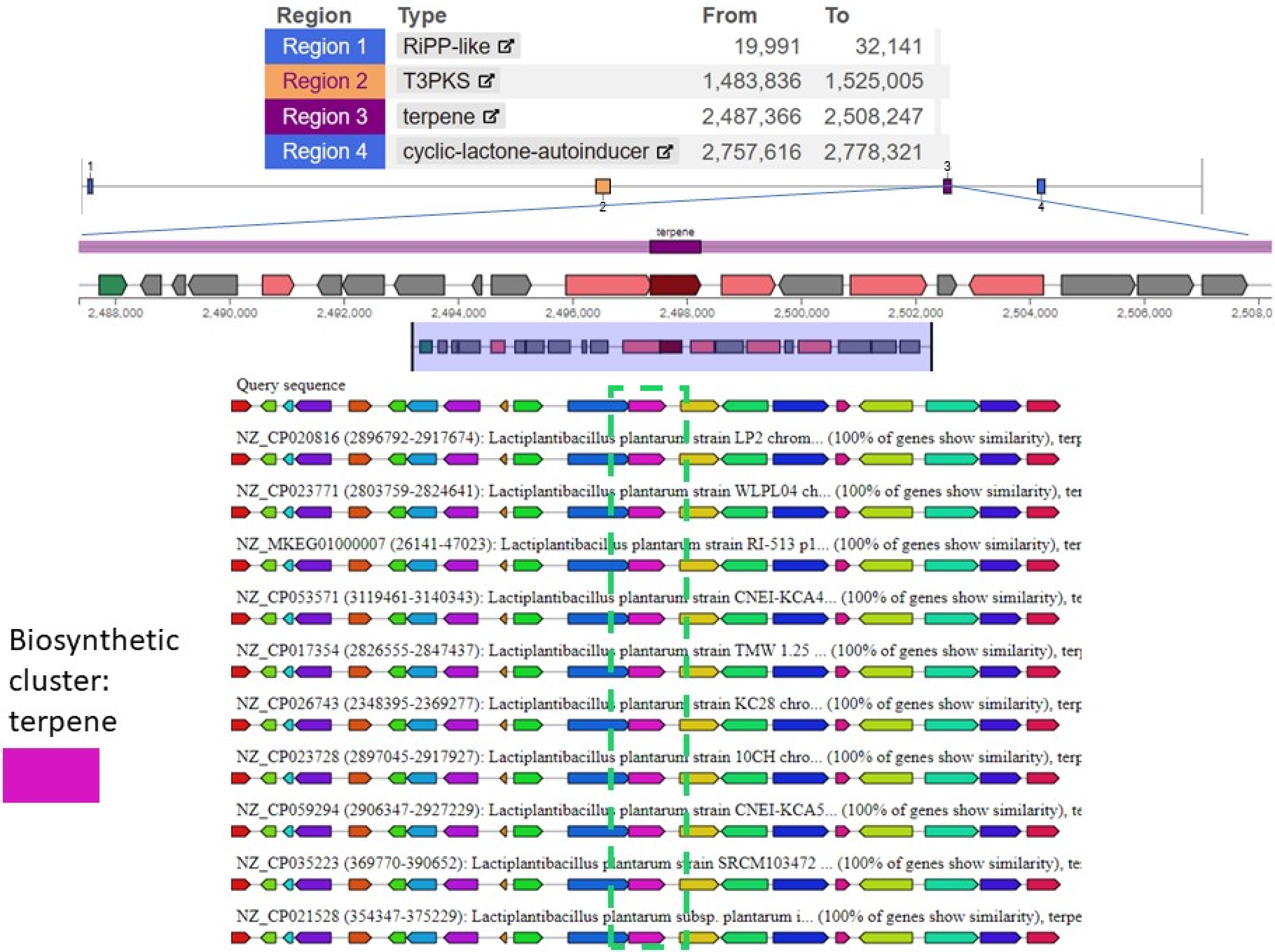
Region | Type | From | To
Region 1 | RiPP-like | 19,991 | 32,141
Region 2 | T3PKS | 1,483,836 | 1,525,005
Region 3 | terpene | 2,487,366 | 2,508,247
Region 4 | cyclic-lactone-autoinducer | 2,757,616 | 2,778,321
terpene
2,488,000
2,490,000
2,492,000
2,494,000
2,496,000
2,498,000
2,500,000
2,502,000
2,504,000
2,506,000
Query sequence
NZ_CP020816 (2896792-2917674): Lactiplantibacillus plantarum strain LP2 chrom... (100% of genes show similarity), terp
NZ_CP023771 (2803759-2824641): Lactiplantibacillus plantarum strain WLPL04 ch... (100% of genes show similarity), ter
NZ_MKEG01000007 (26141-47023): Lactiplantibacillus plantarum strain RI-513 p1... (100% of genes show similarity), ter
NZ_CP053571 (3119461-3140343): Lactiplantibacillus plantarum strain CNEI-KCA4... (100% of genes show similarity), te
NZ_CP017354 (2826555-2847437): Lactiplantibacillus plantarum strain TMW 1.25 ... (100% of genes show similarity), terp
NZ_CP026743 (2348395-2369277): Lactiplantibacillus plantarum strain KC28 chro... (100% of genes show similarity), terp
NZ_CP023728 (2897045-2917927): Lactiplantibacillus plantarum strain 10CH chro... (100% of genes show similarity), terp
NZ_CP059294 (2906347-2927229): Lactiplantibacillus plantarum strain CNEI-KCA5... (100% of genes show similarity), te
NZ_CP035223 (369770-390652): Lactiplantibacillus plantarum strain SRCM103472 ... (100% of genes show similarity), te
NZ_CP021528 (354347-375229): Lactiplantibacillus plantarum subsp. plantarum i... (100% of genes show similarity), terpe
Biosynthetic cluster: terpene

| Region | Type | From | To |
|---|---|---|---|
| Region 1 | RiPP-like | 19,991 | 32,141 |
| Region 2 | T3PKS | 1,483,836 | 1,525,005 |
| Region 3 | terpene | 2,487,366 | 2,508,247 |
| Region 4 | cyclic-lactone-autoinducer | 2,757,616 | 2,778,321 |

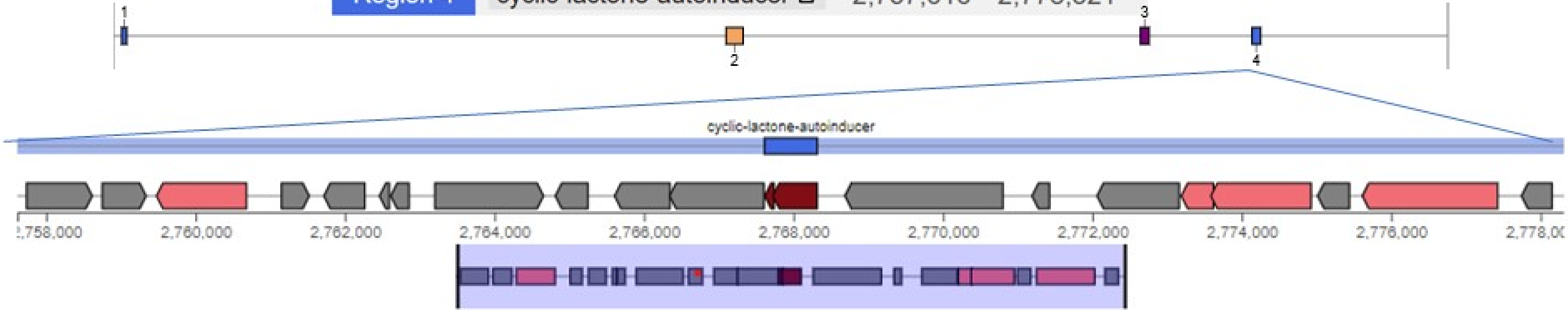
