## Supplementary figures and images for "Exploring the probiotic potential, antioxidant capacity, and healthy aging based on whole genome analysis of *Lactiplantibacillus plantarum* LPJBC5 isolated from fermented milk product"

### Supplementary figure S3

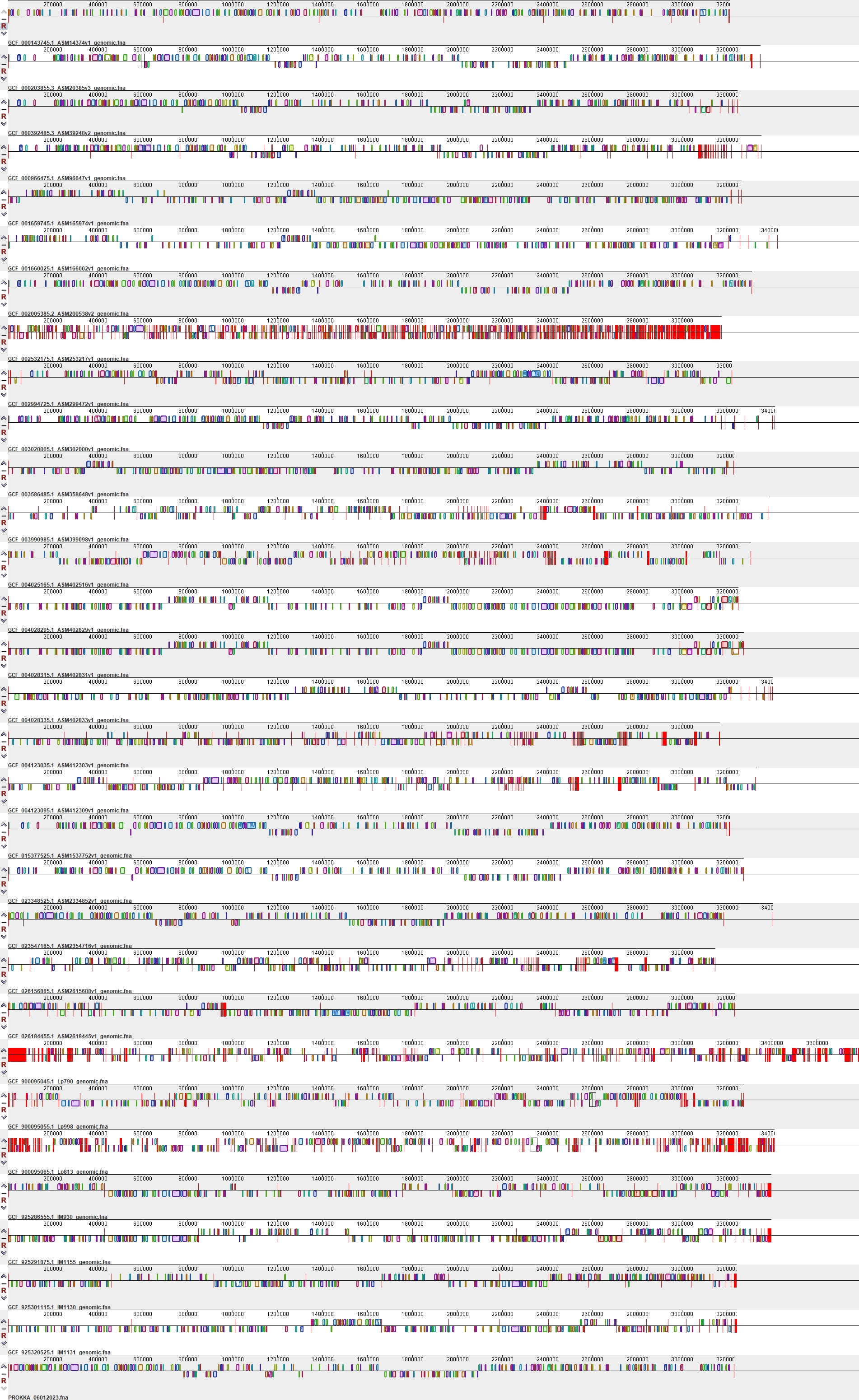
