## Supplementary Table S1 for "Exploring the probiotic potential, antioxidant capacity, and healthy aging based on whole genome analysis of *Lactiplantibacillus plantarum* LPJBC5 isolated from fermented milk product"

**Supplementary Table S1**. Average nucleotide identity (ANI) of *Lactiplantibacillus_plantarum* LPJBC5 genome against existing *Lactobacillus plantarum* genome.

| GENOME1 | GENOME2 | ANI(1->2) | ANI(2->1) |
| --- | --- | --- | --- |
| LPJBC5 | *Lactiplantibacillus_plantarum* ATCC-14917 | 99.71 | 99.71 |
| LPJBC5 | *Lactiplantibacillus_plantarum* CGMCC-1.2437 | 99.74 | 99.74 |
| LPJBC5 | *Lactiplantibacillus_plantarum* DSM-13273 | 99.42 | 99.42 |
| LPJBC5 | *Lactobacillus_plantarum* DSM-16365 | 95.66 | 95.66 |
| LPJBC5 | *Lactobacillus_plantarum* JCM-1149 | 99.72 | 99.72 |
| LPJBC5 | *Lactobacillus_plantarum* NBRC-15891 | 99.71 | 99.71 |
| LPJBC5 | *Lactobacillus_plantarum* NCTC13644 | 99.69 | 99.69 |
