## Supplementary Table S2 for "Exploring the probiotic potential, antioxidant capacity, and healthy aging based on whole genome analysis of *Lactiplantibacillus plantarum* LPJBC5 isolated from fermented milk product"

**Supplementary Table S2**. Comparative genome summary of *Lactiplantibacillus plantarum* strains.

|  | **Strain ID** | **Genome size(mb)** | **G+C content** | **Number of genes** | **Predicted CDS** | **tRNA** | **rRNA** |
| --- | --- | --- | --- | --- | --- | --- | --- |
| *Lactiplantibacillus plantarum* | LPJBC5 | 3.23 | 44.55% | 3098 | 3016 | 65 | 16 |
| *Lactiplantibacillus plantarum* | WCSF1 | 3.3 | 44.42% | 3211 | 3123 | 71 | 16 |
| *Lactiplantibacillus plantarum* | ATCC14917 | 3.21 | 44.50% | 3061 | 2999 | 59 | 2 |
| *Lactiplantibacillus plantarum* | DHCU70 | 3.38 | 44.29% | 3273 | 3195 | 68 | 9 |
| *Lactiplantibacillus plantarum* | 12_3 | 3.4 | 44.44% | 3352 | 3266 | 69 | 16 |
| *Lactiplantibacillus plantarum* | YW11 | 3.24 | 44.52% | 3171 | 3087 | 67 | 16 |
| *Lactiplantibacillus plantarum* | 13_3 | 3.27 | 44.49% | 3208 | 3124 | 67 | 16 |
| *Lactiplantibacillus plantarum* | TK-P2A | 3.21 | 44.64% | 3067 | 2987 | 63 | 16 |
| *Lactiplantibacillus plantarum* | K25 | 3.41 | 44.38% | 3328 | 3244 | 67 | 16 |
| *Lactiplantibacillus plantarum* | LPT52 | 3.41 | 44.38% | 3119 | 3036 | 66 | 16 |
| *Lactiplantibacillus plantarum* | ISO1 | 3.23 | 44.51% | 3081 | 3025 | 52 | 3 |
| *Lactiplantibacillus plantarum* | IM1130 | 3.23 | 44.46% | 3032 | 3104 | 64 | 7 |
| *Lactiplantibacillus plantarum* | IM1131 | 3.24 | 44.46% | 3105 | 3031 | 65 | 8 |
| *Lactiplantibacillus plantarum* | IM1155 | 3.38 | 44.17% | 3232 | 3162 | 62 | 7 |
| *Lactiplantibacillus plantarum* | C4 | 3.22 | 44.53% | 3084 | 3023 | 57 | 3 |
| *Lactiplantibacillus plantarum* | NL42 | 3.35 | 44.42% | 3307 | 3224 | 62 | 16 |
| *Lactiplantibacillus plantarum* | IM930 | 3.39 | 44.17% | 3234 | 3164 | 62 | 7 |
| *Lactiplantibacillus plantarum* | Lp998 | 3.26 | 44.47% | 3172 | 3081 | 74 | 16 |
| *Lactiplantibacillus plantarum* | XZ3303 | 3.32 | 44.41% | 3208 | 3143 | 63 | 1 |
| *Lactiplantibacillus plantarum* | S2.13 | 3.14 | 44.51% | 3058 | 2983 | 65 | 9 |
| *Lactiplantibacillus plantarum* | YW32 | 3.16 | 44.54% | 3041 | 2974 | 63 | 1 |
| *Lactiplantibacillus plantarum* | Lp790 | 3.16 | 44.54% | 3610 | 3524 | 73 | 12 |
| *Lactiplantibacillus plantarum* | Lp813 | 3.36 | 44.42% | 3246 | 3157 | 73 | 15 |
| *Lactiplantibacillus plantarum* | M92C | 3.17 | 44.40% | 2972 | 2917 | 49 | 5 |
| *Lactiplantibacillus plantarum* | Dad-13 | 3.4 | 44.27% | 3337 | 3254 | 66 | 16 |
| *Lactiplantibacillus plantarum* | SKT109 | 3.3 | 44.43% | 3180 | 3120 | 57 | 2 |
| *Lactiplantibacillus plantarum* | P-8 | 3.24 | 44.55% | 3180 | 3094 | 69 | 1 |
| *Lactiplantibacillus plantarum* | LZ227 | 3.42 | 44.35% | 3386 | 3297 | 72 | 16 |
| *Lactiplantibacillus plantarum* | LZ206 | 3.26 | 44.55% | 3250 | 3166 | 67 | 16 |
| *Lactiplantibacillus plantarum* | 10CH | 3.31 | 44.51% | 3157 | 3074 | 66 | 16 |
