## Supplementary Table S3 for "Exploring the probiotic potential, antioxidant capacity, and healthy aging based on whole genome analysis of *Lactiplantibacillus plantarum* LPJBC5 isolated from fermented milk product"

**Supplementary Table** S**3**. Distribution of core, accessory, unique, and exclusively absent genes among 31 *Lactiplantibacillus plantarum* strains

| **Sl no.** | **Organism name** | **No. of core genes** | **No. of accessory genes** | **No. of unique genes** | **No. of exclusively absent genes** |
| --- | --- | --- | --- | --- | --- |
| 1 | *Lactiplantibacillus plantarum* ATCC14917 | 1844 | 911 | 5 | 4 |
| 2 | *Lactiplantibacillus plantarum* WCSF1 | 1844 | 943 | 51 | 9 |
| 3 | *Lactiplantibacillus plantarum* P-8 | 1844 | 830 | 6 | 0 |
| 4 | *Lactiplantibacillus plantarum* NL-42 | 1844 | 883 | 64 | 23 |
| 5 | *Lactiplantibacillus plantarum* LZ206 | 1844 | 865 | 27 | 34 |
| 6 | *Lactiplantibacillus plantarum* LZ227 | 1844 | 919 | 66 | 12 |
| 7 | *Lactiplantibacillus plantarum* 10CH | 1844 | 927 | 25 | 1 |
| 8 | *Lactiplantibacillus plantarum* M92C | 1844 | 892 | 6 | 42 |
| 9 | *Lactiplantibacillus plantarum* C4 | 1844 | 914 | 0 | 4 |
| 10 | *Lactiplantibacillus plantarum* K25 | 1844 | 907 | 69 | 4 |
| 11 | *Lactiplantibacillus plantarum* DR7 | 1844 | 873 | 42 | 2 |
| 12 | *Lactiplantibacillus plantarum* DHCU70 | 1844 | 1010 | 77 | 4 |
| 13 | *Lactiplantibacillus plantarum* SKT109 | 1844 | 1003 | 2 | 0 |
| 14 | *Lactiplantibacillus plantarum* YW11 | 1844 | 799 | 0 | 0 |
| 15 | *Lactiplantibacillus plantarum* 13_3 | 1844 | 801 | 0 | 7 |
| 16 | *Lactiplantibacillus plantarum* 12_3 | 1844 | 989 | 45 | 5 |
| 17 | *Lactiplantibacillus plantarum* YW32 | 1844 | 849 | 3 | 1 |
| 18 | *Lactiplantibacillus plantarum* XZ3303 | 1844 | 997 | 7 | 0 |
| 19 | *Lactiplantibacillus plantarum* TK-P2A | 1844 | 901 | 0 | 7 |
| 20 | *Lactiplantibacillus plantarum* LPT52 | 1844 | 883 | 59 | 2 |
| 21 | *Lactiplantibacillus plantarum* Dad-13 | 1844 | 718 | 104 | 43 |
| 22 | *Lactiplantibacillus plantarum* S2.13 | 1844 | 821 | 29 | 20 |
| 23 | *Lactiplantibacillus plantarum* IS01 | 1844 | 860 | 1 | 4 |
| 24 | *Lactiplantibacillus plantarum* Lp790 | 1844 | 1057 | 70 | 15 |
| 25 | *Lactiplantibacillus plantarum* Lp998 | 1844 | 945 | 1 | 0 |
| 26 | *Lactiplantibacillus plantarum* Lp813 | 1844 | 969 | 34 | 2 |
| 27 | *Lactiplantibacillus plantarum* IM930 | 1844 | 1060 | 0 | 0 |
| 28 | *Lactiplantibacillus plantarum* IM1155 | 1844 | 1070 | 7 | 0 |
| 29 | *Lactiplantibacillus plantarum* IM1130 | 1844 | 923 | 1 | 0 |
| 30 | *Lactiplantibacillus plantarum* IM1131 | 1844 | 925 | 0 | 0 |
| 31 | *Lactiplantibacillus plantarum* LPJBC5 | 1844 | 893 | 45 | 7 |
